# Peptide-Specific CARs Recognize WT1 Promiscuously Presented by Diverse HLA Class II Alleles

**DOI:** 10.1101/2025.06.30.660492

**Authors:** Chung-Hsi Wang, Evey Y. F. Zheng, Toshiki Ochi, Yota Ohashi, Fumie Ihara, Saori Fukao, Giselle M Boukhaled, Ben X. Wang, Dong-Hoon Han, Xinyu Wei, Brian D. Burt, Kayoko Saso, Yukiko Matsunaga, Dalam Ly, Yuki Kagoya, Marcus O. Butler, Mark D. Minden, Naoto Hirano

**Author notes:** These authors contributed equally to this work (Chung-Hsi Wang, Evey Y. F. Zheng). Correspondence and requests for materials should be addressed to Naoto Hirano, MD, PhD., Naoto Hirano, MD, PhD, Princess Margaret Cancer Centre, 610 University Avenue, Toronto, ON M5G 2M9, Canada.

## Abstract

Chimeric antigen receptor (CAR) technology has revolutionized B-cell malignancy treatment by enabling T cells to effectively recognize and target cancer-specific surface antigens. However, CAR T cells show limited efficacy against other blood cancers and solid tumors due to challenges in identifying suitable surface targets. Here, we present a novel approach to CAR development, targeting the intracellular Wilms’ tumor 1 (WT1) oncoprotein, cross-presented by surface HLA-class II (HLA-II) alleles. WT1-CAR T cells, derived from an antibody raised solely against a WT1 peptide, recognized the WT1330-348 peptide promiscuously presented by 18 out of 20 tested HLA-II alleles, overcoming traditional HLA restrictions. WT1-CAR T cells specifically recognized leukemic cells in a WT1 and HLA-II-dependent manner and mediated an antitumor response in vitro and in vivo. This innovative approach to CAR T cell development transcends traditional HLA restrictions and offers a promising therapeutic option to a wide and genetically diverse patient population.

**Statement of significance:** This study describes a novel CAR T therapy approach leveraging the distinctive and shared characteristic of HLA-II-peptide binding promiscuity, enabling targeting of the intracellular oncoprotein WT1 presented across diverse HLA-II families. Our study demonstrates a viable framework for designing CAR T therapies that benefit genetically diverse patient populations.

## Introduction

T cell receptors (TCRs) and TCR-like chimeric antigen receptors (CARs) targeting human leukocyte antigen (HLA)-restricted tumor-associated antigens (TAAs) and neoantigens represent promising immunotherapeutic strategies(1–4). To date, the majority of antitumor TCR and TCR-like CARs in development intuitively target peptides presented by HLA-class I (HLA-I) molecules, which serve as the canonical machinery for endogenous antigen presentation. However, the clinical safety of HLA-I-targeting therapies is threatened by “on-target, off-tumor” toxicity resulting from the ubiquity of HLA-I molecules (5).

In contrast, HLA-class II (HLA-II) molecules exhibit more restricted expression, typically limited to professional antigen-presenting cells (APCs), thymic epithelial cells (TECs), and activated T cells. In addition to the canonical presentation of peptides derived from exogenous proteins, HLA-II can also present endogenously-derived peptides, mediated by transporter associated with antigen processing (TAP) protein and autophagy (6,7). Notably, common HLA-II molecules, such as HLA-DP2 and DP4, have also been shown to constitutively present peptides derived from both exogenous and endogenous sources to CD4^+^ T cells (8,9). Moreover, various malignancies have been reported to aberrantly upregulate HLA-II expression, further supporting its potential as a tumor-selective target (10). These observations highlight the underexplored therapeutic potential of targeting HLA-II-presented antigenic peptides in cancer immunotherapy, particularly in light of their capacity to present intracellular tumor-associated antigens with limited expression in healthy tissues.

In addition, a major limitation to the broad application of currently available T cell therapies lies within the therapeutic specificity to a single peptide-HLA complex. Although a few TCRs have been identified with restriction to multiple HLAs, this is typically confined to alleles within the same family (11–13). Yet, modalities that recognize a single antigen across diverse HLA superfamilies are not unheard of. A notable precedent is CerCLIP.1, initially developed to detect the class II-associated invariant chain peptide (CLIP) presented by DR molecules, has since been used to detect CLIP complexed with HLA-II molecules across DP, DQ and DR families (8,14–16). Importantly, CerCLIP.1 was raised against the CLIP peptide alone, independent of complexation with any HLA-II molecules (14). This observation suggests that by leveraging the promiscuous binding of peptides to HLA-II molecules, it is possible to develop modalities that recognize a common peptide presented across different HLA-II alleles.

As a proof of concept, we employed such a strategy to generate monoclonal antibodies (mAbs) specific to peptides derived from Wilms’ tumor 1 (WT1) protein, a well-established TAA with a favorable safety profile and a long-standing history as a target in various cancer immunotherapeutic strategies (17,18).

## Results

### Generation of a CAR targeting WT1 presented across multiple HLA-II allotypes

Previous studies have reported that WT1_332-348_ peptide exhibits the capacity to bind HLA-II alleles across multiple HLA-II supertypes and elicit TCR-mediated immune responses (19–22). To generate a CAR capable of targeting WT1 presented across HLA-II alleles, we immunized mice with WT1_330-348_, a longer peptide with a cysteine residue at position 330 that is critical for N-terminal conjugation with adjuvant for enhanced immunogenicity. Following immunization with the peptide, multiple hybridoma clones specific to WT1_330-348_ were generated and assessed for their reactivity against the unbound peptide and the peptide in complex with HLA-DP4. Among these clones, clone 5H2 exhibited the ability to bind the WT1_330-348_ peptide alone and when presented in the context of HLA-DP4 (HLA-DP4/WT1_329-348_, Supplementary Fig. S1A). Bio-layer interferometry (BLI) analysis further revealed that clone 5H2 bound to WT1_330-348_ peptide with an affinity of K_D_ ≈50 nM, and to the HLA-DP4/WT1_329-348_ complex with a K_D_ ≈13 μM (Supplementary Fig. S1B-C). Notably, this latter value falls within the intermediate micromolar affinity range (10–100 μM) characteristic of TCRs recognizing peptide–HLA (pHLA) complexes. Affinities in this range are considered biologically and therapeutically optimal, as they enable selective recognition of TAAs while minimizing off-tumor reactivity (23–25). Based on these findings, clone 5H2 was utilized to develop a second-generation CAR, designated as WT1-CAR (Fig.1A) (26).

**Figure 1:**
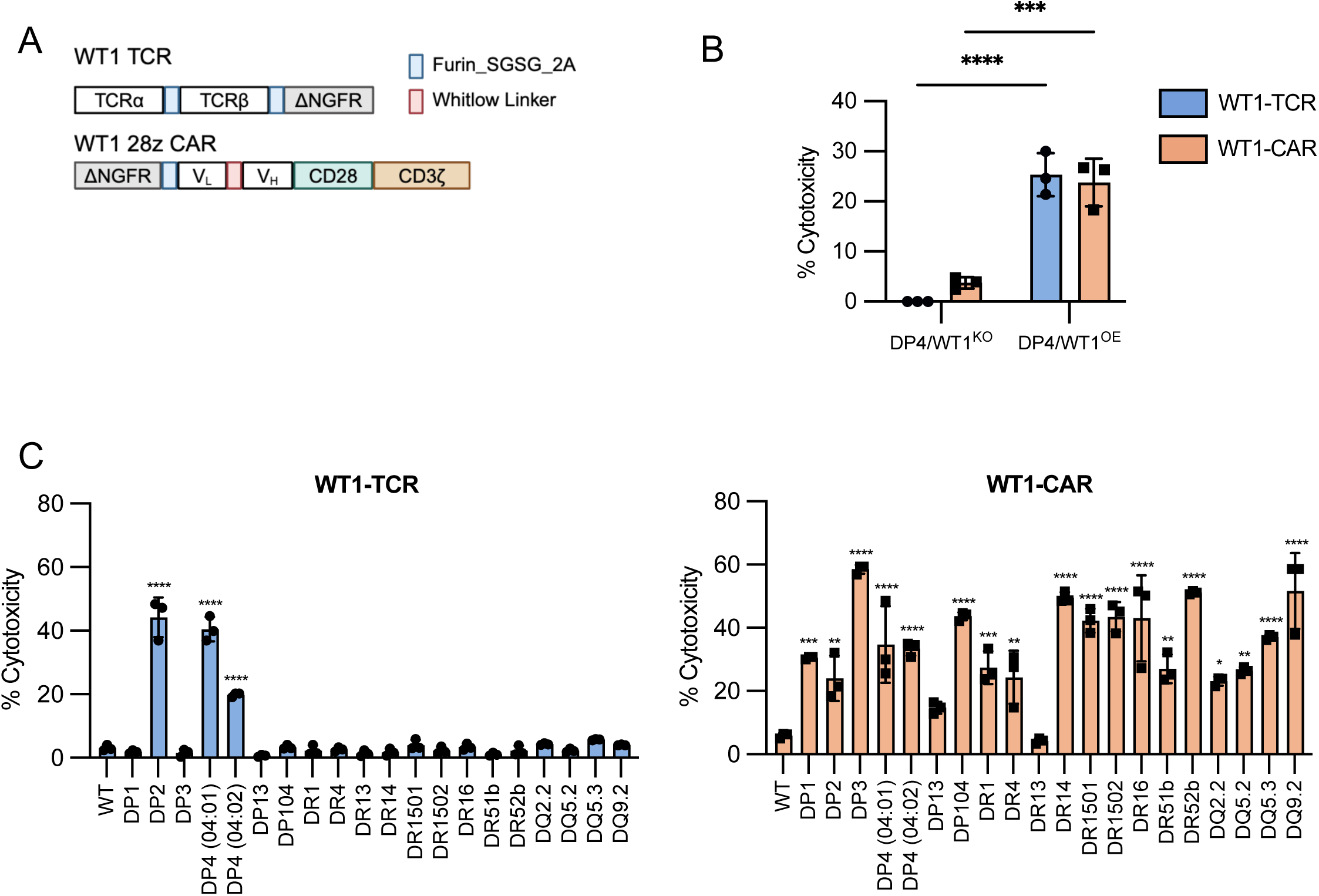
Establishing CAR T cells targeting WT1 peptide presented by HLA-DP4 and other HLA-II molecules. (A) Schematic representation of the WT1-TCR and WT1-CAR constructs. (B) Cytotoxicity of WT1-TCR or WT1-CAR T cells against the indicated K562 expressing DP4 with WT1 gene knockout (KO) or overexpression (OE) at an E:T ratio of 5:1 for 18 h, measured by flow cytometry-based killing assay. n = 3 donors. (C) Cytotoxicity of WT1-TCR or -CAR T cells against WT1^OE^ K562 expressing the indicated HLA-II molecules at an E:T ratio of 5:1 for 18 h was measured by flow cytometry-based killing assays. HLA-II-null, WT1^OE^ K562 (WT) was used as a negative control. n = 3 donors. (B-C) Data shown represents the mean ± SD of all donors. *P<0.05, **P<0.01, ***P<0.001, ****P<0.0001 by one-way ANOVA multiple comparison test with Bonferroni tests.

To gain insight into how the newly generated WT1-CAR recognizes WT1 compared to conventional TCRs, we incorporated a previously characterized WT1-specific TCR, termed WT1-TCR, confirmed to be able to recognize WT1_330-348_ presented in the context of HLA-DP2,-DP4, and -DP5 (Supplementary Fig. S1D-E) (21). Target cells generated from HLA-II-deficient K562 cells, engineered to overexpress full-length WT1 (K562/WT1^OE^) or have endogenous WT1 gene deletion (K562/WT1^KO^, Supplementary Fig. S1F), were transduced to express HLA-DP4.

Consistent with previous observations, WT1-TCR transduced T cells (WT1-TCR T cells) showed recognition and cytolysis of K562/DP4/WT1^OE^ (DP4/WT1^OE^) but not K562/DP4/WT1^KO^ (DP4/WT1^KO^, Fig. 1B). Similarly, WT1-CAR T cells showed analogous recognition, including cytolysis of DP4/WT1^OE^ but not DP4/WT1^KO^ (Fig. 1B), suggesting that both WT1-TCR and the newly generated WT1-CAR could recognize a naturally processed WT1 epitope presented in the context of HLA-DP4.

Considering the potential for WT1_330-348_ presentation across diverse HLA-II alleles and the design of our WT1 targeting CAR construct, we explored the possibility that WT1-CAR could recognize WT1 presented in the context of different HLA-II allotypes. To investigate the recognition of endogenously processed WT1, and to control for potential differences in HLA-II affinity for WT1 peptides, we generated K562/WT1^OE^ cells individually expressing 20 different HLA-II molecules. Despite the ectopic overexpression of WT1, WT1-TCR T cells induced cytotoxicity against only HLA-DP2 and -DP4 expressing target cells, retaining their HLA restriction (Fig. 1C). In contrast, WT1-CAR demonstrated cytotoxic activity against K562/WT1^OE^ co-expressing 18 out of 20 tested HLA-II alleles, including six HLA-DP, eight HLA-DR, and four HLA-DQ alleles (Fig. 1C), but not HLA-II negative K562/WT1^OE^ cells.

### WT1-CAR shows N-terminal-biased recognition of the broadly presented WT1_330-348_ peptide

Given the capacity of WT1-CAR to recognize WT1 presented across multiple HLA-II alleles, we sought to define the minimal epitope required for recognition. We hypothesized that while both WT1-TCR and WT1-CAR receptors necessitate WT1 peptide presentation within the context of the HLA-II binding groove, the WT1-CAR, since being generated against a free peptide, may exhibit binding to residues beyond interactions that incorporate the HLA-II molecule. To investigate these possibilities, we performed a series of assays using truncated peptides and alanine scanning mutagenesis to map the minimal epitope required for WT1-TCR and -CAR recognition.

We first compared the minimal epitope required for TCR and CAR recognition of the HLA-DP-restricted WT1. HLA-II deficient T2 cells expressing HLA-DP2, -DP4, or -DP5 were pulsed with a panel of sequentially truncated WT1_328-348_ peptides at the N- and/or C-terminal ends and their abilities to elicit T cell responses were assessed. Despite the deletion of N-terminal residues 328-333(_328_PGCNKR_333_), HLA-DP presentation of truncated peptides was tolerated, as confirmed by the preserved WT1-TCR-mediated responses (Fig. 2A and Supplementary Fig. S2A-B, left).

**Figure 2:**
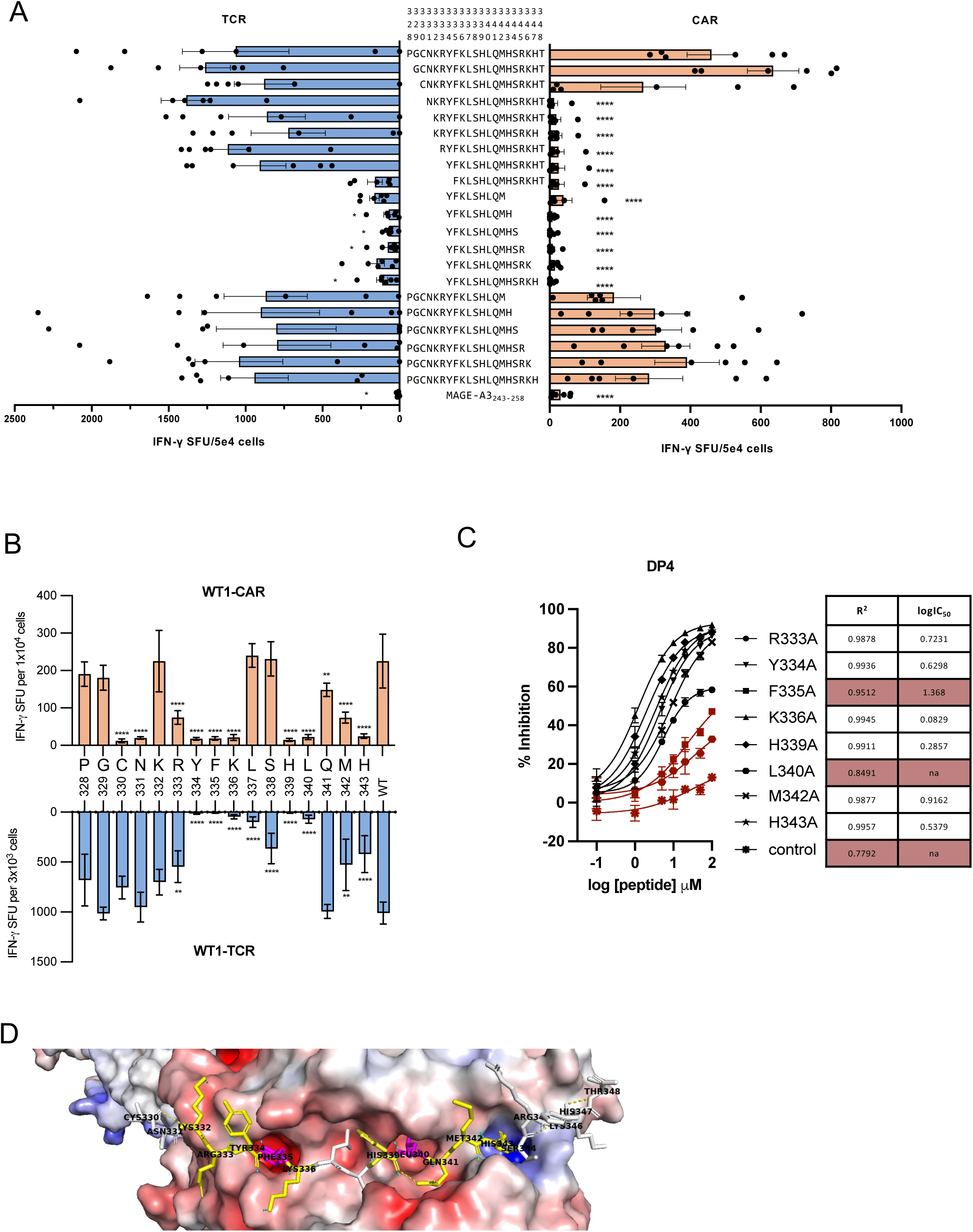
WT1-TCR and CAR have distinct modes of recognition of WT1 peptide presented with a homologous binding motif across HLA-II. (A) WT1-TCR or CAR T cells were stimulated with T2/DP4 pulsed with different deletion peptides for 20-24 h. MAGE-A3_243-258_ was used as a negative control. IFN-γ secretion was measured by ELISpot. n = 6 donors. *P<0.05, **P<0.01, ***P<0.001, ****P<0.0001 by one-way ANOVA multiple comparison test with Bonferroni tests. (B) WT1-TCR or CAR T cells were stimulated with T2/DP4 pulsed with WT1_328-348_ (WT) or WT1_328-348_ peptide with alanine substitution at indicated amino acid positions for 20-24 h. IFN-γ secretion was measured by ELISpot analysis. n = 3 donors. (A-B) Data showing the mean ± SD of all donors. (C) Competitive binding assays with T2 cells expressing HLA-DP4, pulsed with graded concentrations of WT1_328-348_ alanine mutant at specified positions in the presence of 1 µM biotinylated-CLIP peptide. Poor binders (Methods) are marked in red. Data showing the mean ± SD from three independent studies. (D) Structural modelling of WT1_330-348_ presented in HLA-DP4 by AlphaFold3 showing HLA surface electrostatic charge (red: negative, blue: positive), peptide-HLA H-bonds (light blue dash), intra-peptide H-bonds (yellow dash), anchors identified in 2B (magenta), residues forming H-bonds (yellow).

Further analysis showed that peptides lacking N-terminus (_334_YFKLSHLQMHSRKHT_348_) or C-terminal (_328_PGCNKRYFKLSHLQM_342_) residues retained the ability to activate WT1-TCR T cells, identifying an overlapping core region consisting of _334_YFKLSHLQM_342_. Though this overlapping peptide alone failed to elicit WT1-TCR T cell response, requiring additional N- and C-terminal flanking residues, this region likely represents the core HLA-DP4 peptide binding region (Fig. 2A and Supplementary Fig. S2A, left). In contrast, WT1-CAR T cell activation was abrogated upon truncation of N-terminal residues 328-330 (_328_PGC_330_), suggesting that WT1-CAR recognized additional distal N-terminal residues beyond those needed by WT1-TCR, including but not limited to C330 (Fig. 2A, right). In support of these observations, deletion of the C-terminal residues 344-348 (_344_HSRKH_348_) did not impact the recognition by either WT1-CAR or -TCR T cells (Fig. 2A).

Presentation of truncated peptides by HLA-DP2 produced highly comparable results, supporting the conservation of peptide presentation across these two alleles (Supplementary Fig. S2A). However, when tested with T2 cells expressing HLA-DP5, the T cell response profile displayed slight deviations. In particular, the overlapping segment required for successful peptide loading was defined as _332_KRYFKLSHLQMH_343_, suggesting subtle differences in peptide register and binding preferences across HLA-DP alleles (Supplementary Fig. S2B). Collectively, across all HLA-DPs tested, deletion of the N-terminal residues, particularly C330, but not C-terminal residues, abrogated WT1-CAR T cell recognition, reinforcing the observation that WT1-CAR engagement is uniquely dependent on N-terminal “jagged ends”, distal to the HLA-binding groove, despite the slightly shifted peptide presentation across different alleles.

To further determine the individual residues responsible for receptor-recognition, we performed alanine scanning mutagenesis within the N-terminal jagged end and the binding core for their ability to stimulate WT1-CAR or WT1-TCR transduced T cells in the context of HLA-DP4. Consistent with N-terminal truncated peptides, L337 and S338 residues were required only by WT1-TCR T cell for recognition, while WT1-CAR T recognition was uniquely dependent on C330, N331 (Fig. 2B). Additionally, mutations within the proposed HLA-DP4 peptide binding core at positions R333, Y334, F335, K336, H339, L340, M342, and H343 significantly impaired recognition by both receptors. To evaluate whether this pattern was conserved in other HLA-II alleles, we extended the alanine scanning analysis to representative members of other HLA-II families. In target cells expressing HLA-DQ9.2 or -DR1501, WT1-TCR and -CAR exhibited nearly identical residue dependency profiles to that observed with HLA-DP4 (Supplementary Fig. S3A).

### Presentation of WT1_330-348_ by HLA-II alleles involves homologous core binding motifs and distinct anchor residues

While mutation analysis of WT1_328-343_ identified shared and distinct residues required for recognition by WT1-TCR and WT1-CAR, we further examined the importance of residues in HLA-II binding and presentation using series of cell-free competitive peptide binding assays. HLA-binding capabilities of mutant peptides were evaluated relative to the invariant chain CLIP peptide. For HLA-DP4, mutations at positions F335 and L340, exhibited substantially reduced competitive capacity—as measured by IC50 value or dose-inhibition model fitting (R^2^<0.9)— indicating that F335 and L340 are critical for peptide binding to HLA-DP4 (Fig. 2C). We then employed AlphaFold3 to examine the peptide conformation and interaction with HLA anchor pockets (27). Notably, F335 and L340 are positioned within the deep negatively charged P1 and P6 anchor pockets, respectively (Fig. 2D). Neighbouring regions —_332_KR(Y)FL_336_, and _339_H(L)QMH_343_ — form hydrogen bonds with HLA chains, likely contributing to peptide stability and presentation. Though not primary anchors, mutations in these surrounding residues influence receptor recognition, as seen by the mutagenesis analyses (Fig. 2B).

To extend the analysis to HLA-DR and HLA-DQ molecules, we evaluated residue contributions across HLA-DR1501 and -DQ9.2. For HLA-DR1501, residues R333, Y334, L340, and M342 were crucial for binding (Supplementary Fig. S3B). Taken together with AlphaFold3 modelling, L340 and M342 correspond to P4 and P6 anchor positions, forming anchoring stabilizing contacts. Additional interactions were observed in _335_FKLS_338_ and _334_SR_345_, forming hydrogen bonds around the P1 and P9 pockets. Noticeably, the N-terminal flanking region forms multiple intra-peptide H-bonds between N331, K332, R333 and Y334 (Supplementary Fig. S3C). In HLA-DQ9.2, key anchors were identified at H339 and M342, with the segment _333_RYFK_336_ appearing particularly sensitive to mutation, suggesting it may contain additional anchoring residues or be structurally flexible and making it susceptible to conformational changes that impact peptide presentation (Supplementary Fig. S3D). Overall, despite some variability in register usage, competitive binding assays and *in silico* analyses revealed a highly homologous core binding motif across HLA-DP4, -DQ9.2 and -DR1501 where the essential anchor residues-- positioned within _334_YF_335_ and _340_LQM_342_ -- are largely conserved.

### WT1-CAR T cells effectively recognize WT1 presented by multiallelic HLA-II targets

With the unique ability of WT1-CAR to recognize WT1 presented across divergent HLA-II alleles, we sought to mimic natural settings, where cells co-expressing multiple HLA-II genes may have allele-specific bias in WT1 presentation (28,29). To explore this possibility, we generated K562/WT^OE^ co-expressing HLA-DP4, -DR1501, and -DQ9.2 (Supplementary Fig. S4A) and compared their stimulatory capacity to K562 cells expressing HLA-II molecules individually. The reactivity of WT1-CAR T cells was measured over 85 h using the xCelligence Real-Time Cell Analysis (RTCA) system. WT1-CAR T cells demonstrated significantly higher cytotoxicity against HLA-DQ9.2 target cells compared to HLA-DR1501 or -DP4 target cells (Fig.3A). Notably, WT1-CAR T cell cytotoxicity towards multiallelic HLA-II target cells was superior to that of monoallelic HLA-DR1501 and -DP4 target cells, but was not statistically significant compared to that of monoallelic HLA-DQ9.2 target cells (Fig. 3A). Nonetheless, these results suggest that in the presence of multiple HLA-II alleles, WT1 may be presented by the majority of these alleles, if not all, albeit with variable presentation efficiencies.

**Figure 3:**
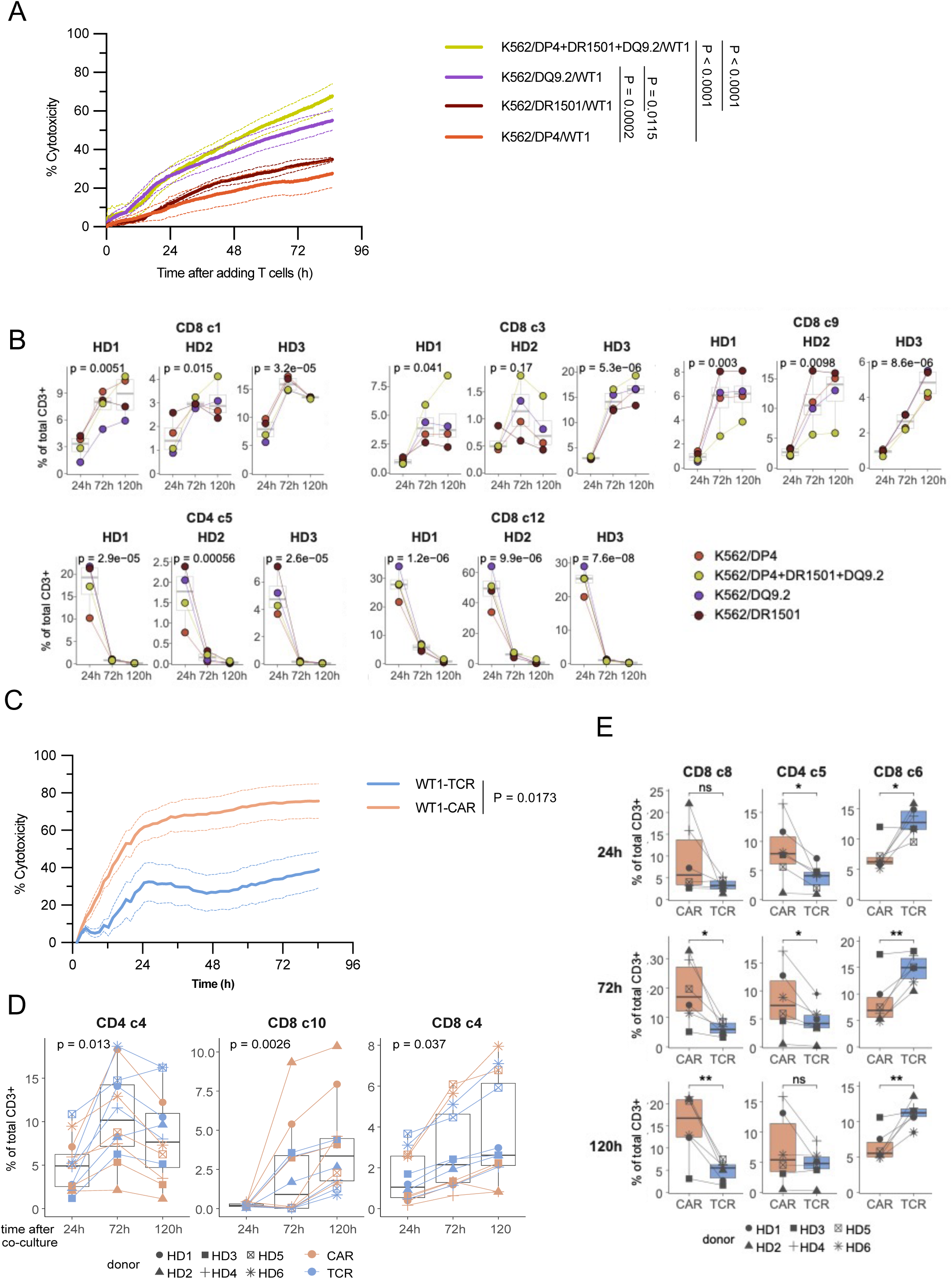
WT1-CAR T cells effectively recognize target cells simultaneously presenting WT1 peptide on multiple HLA-II molecules. (A) Long-term cytotoxicity of WT1-CAR T cells against WT1^OE^ K562 expressing the indicated HLA-II molecules at an E:T ratio of 1:5 was measured by impedance-based xCELLigence killing assays. Target cell cytotoxicity was calculated based on cell index (Methods). n = 5 donors. Data showing the mean ± SD of all donors. P values calculated at endpoint (85 h) with two-way ANOVA multiple comparison test with Tukey tests. (B) Cell cluster populations (>3%) that were identified to be significantly influenced by time in all T cells as determined by two-way paired ANOVA. (C) Long-term cytotoxicity of WT1-TCR or CAR T cells against target cells K562/DP4+DQ1502+DR9.1/WT1 at an E:T ratio of 20:1 measured by fluorescence signal-based xCELLigence killing assay. Target cell cytotoxicity was calculated based on total fluorescence intensity (Methods). n = 5 donors. Data showing the mean ± SD of all donors. P value calculated with cytotoxicity value at 85 h with paired t-test. (D) Cell cluster populations (>2%) that were identified to be significantly influenced by time as determined using two-way paired ANOVA. (E) Cell cluster populations (>5%) that were identified to be significantly influenced by effector cell type as determined using two-way paired ANOVA followed by t-test comparison of each day.

To further analyze the phenotypic changes that may occur in WT1-CAR T cells stimulated by monoallelic or multiallelic targets, we used a 36-marker high-dimensional mass cytometry (CyTOF) panel. Dimensionality reduction and unsupervised clustering on pooled data from WT1-CAR T cells (CD45^+^CD3^+^ cells; Supplementary Fig. S4B) activated by monoallelic or multiallelic target cells across 120 h revealed 13 unique CD8^+^ T cell clusters (CD8 c1-c13), five CD4^+^ T clusters (CD4 c1-c5) and two CD4^-^CD8^-^ clusters (DN1-2) (Supplementary Fig. S4C). No statistically significant difference was observed between T cells stimulated by monoallelic or multiallelic targets at any time point. However, when analyzed across time points, we observed the enrichment and reduction of clusters (Supplementary Fig. S4D). CD8 c1, c3, and c9, consisting of GzmB^+^perforin^+^ cells, were observed to expand, while CD8 c12 and CD4 c5, consisting of early activated/naïve CD45RA^+^ T cells rapidly diminished over time (Fig. 3B and Supplementary Fig. S4E). These results suggest that WT1-CAR T cells showed a similar activation phenotype regardless of the expression breadth of HLA-II alleles.

To determine whether differential receptor recognition of WT1 influences the functional and phenotypic properties of T cells, we compared the cytotoxic capacity of WT1-CAR and WT1-TCR T cells against multiallelic K562/WT^OE^ target cells. WT1-CAR T cells exhibited significantly higher cytotoxicity than WT1-TCR T cells after 85 h of co-culture (Fig. 3C). The phenotypic differences between transduced T cells were analyzed using CyTOF analysis of the co-cultured T cells. Dimensionality reduction and unsupervised clustering revealed seven CD4^+^ T cell clusters (CD4 c1-c7), 12 CD8^+^ T cell clusters (CD8 c1-c12), three double negative clusters (DN 1-3) and one double positive cluster (DP 1) with distinct distributions of WT1-CAR and WT1-TCR T cells across these subsets (Supplementary Fig. S5A-B). CD4 c4 and CD8 c10, consisting of proliferating Ki67^+^ cells and CD8 c4, consisting of GzmB^+^perforin^+^ T cells, expanded irrespective of the transduced receptor, suggesting that target cells successfully induced T cell responses in both WT1-TCR and WT1-CAR T cells (Fig. 3D and Supplementary Fig. S5C). CD8 c8 (CD45RA^+^CCR7^-^ perforin^+^GzmB^+^PD1^+^) and CD4 c5 (CD45RO^+^FoxP3^+^) T cells were consistently more enriched in WT1-CAR T cells compared to WT1-TCR, whereas WT1-TCR T cells were more prevalent in cluster CD8 c6 (CD8^+^CD45RA^+^CCR7^+^) compared to WT1-CAR T cells (Fig. 3E and Supplementary Fig. S5C). These results indicate that WT1-CAR T cells experienced higher levels of antigen activation compared to WT1-TCR T cells. This difference may be attributed to the broader availability of antigen on target cell surfaces, as WT1-TCR recognizes a single pHLA-II allele, whereas WT1-CAR recognizes three distinct HLA-II-presented WT1 peptides.

### WT1-CAR T cells demonstrate anti-AML activity with minimal on-target and off-target toxicities *in vitro*

WT1 belongs to the highly homologous family of zinc-finger nucleases (30). To assess the potential cross-reactivity of WT1-CAR T cells towards analogous peptides, we identified candidate peptides with sequence homology to WT1_330-348_ using BLAST (Methods, Supplementary Table S1). T2 cells individually expressing HLA-DP4, -DR1501, or -DQ9.2 were pulsed with WT1_330-348_ or each of the analogous peptides and co-cultured with WT1-CAR T cells. WT1-CAR T cells did not show detectable reactivity to any candidate peptide (Supplementary Fig. S6A). Notably, all the candidate peptides studied included C330, one of the key residues for WT1-CAR recognition (Fig.2 and Supplementary Table S1), suggesting these analogous peptides are not presented by HLA-II in a similar conformation that exposes the N-terminal end for CAR recognition. Additionally, as WT1 contributes to blood cell development and is expressed in early hematopoietic precursors (31), we assessed the potential “on-target, off-tumor toxicity” of the WT1-CAR T cells on CD34^+^ hematopoietic stem and progenitor cells (HSPCs) that are HLA-II^+^/WT1^+^ (Supplementary Fig. S6B). The *in vitro* co-culture assays did not detect WT1-CAR T cell-dependent cytotoxicity against CD34^+^ HSPCs, indicating minimal recognition of HLA-II/WT1 complexes on normal CD34^+^ HPCs (Fig. 4A). Collectively, these results suggest that though WT1-CAR T cells promiscuously recognize HLA-II/WT1, specificity toward the WT1 epitope is strictly retained.

**Figure 4:**
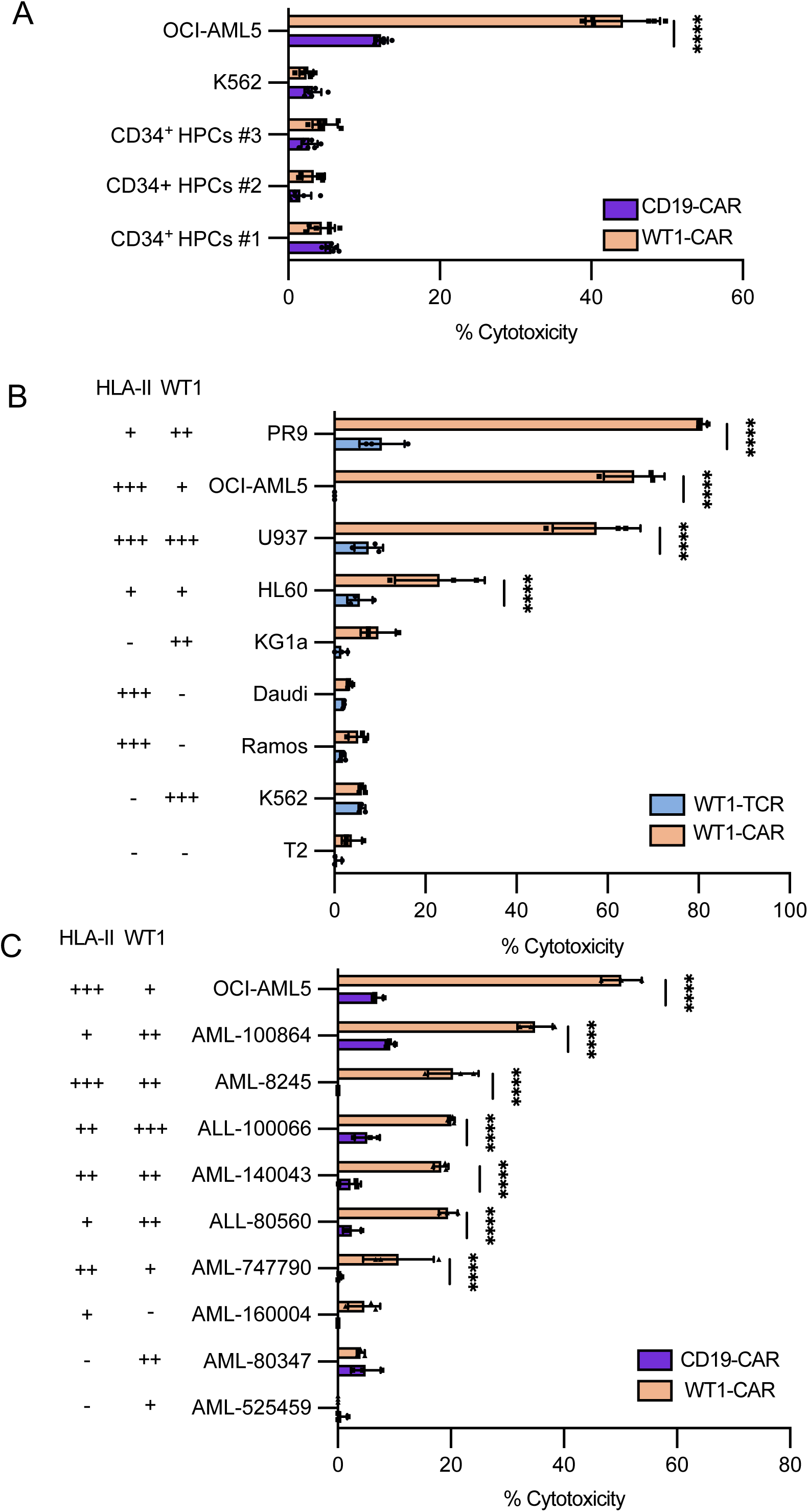
WT1-CAR T cells recognize various WT1^+^/HLA-II^+^ leukemia and lymphoma cell lines and primary leukemic samples without “on-target, off-tumor” toxicities. (A) Cytotoxicity of WT1-CAR or control-CAR (CD19-specific) T cells against indicated cell lines and human primary CD34^+^ hematopoietic cells at an E:T ratio of 5:1 for 6 h as measured by flow cytometry-based killing assays. n = 6 donors. (B) Cytotoxicity of WT1-TCR or CAR T cells against the indicated leukemia and lymphoma cell lines at an E:T ratio of 1:1 for 18 h was measured by flow cytometry-based killing assays. n = 3 donors. (C) Cytotoxicity of WT1-CAR or CD19-CAR (control) transduced T cells against primary ALL or AML samples at an E:T ratio of 5:1 for 18 h was measured by flow cytometry-based killing assays. n = 3 donors. (B-C) HLA-II and WT1 expression were measured by flow cytometric analysis. < 3% positive (-); 3-33% positive (+); 33-66% positive (++); > 66% positive (+++). (A-C) Data shown represents the mean ± SD of all donors. ****P<0.0001 by two-way ANOVA with Bonferroni tests.

We next investigated the ability of WT1-CAR T cells to target blood cancer cell lines endogenously expressing HLA-II and WT1 (Supplementary Fig. S7A). WT1-CAR T cells demonstrated potent HLA-II/WT1 dependent cytotoxicity towards HL-60, PR9, OCI-AML5, and U937 cell lines but not towards cell lines lacking either WT1 and/or HLA-II expression (Fig. 4B). Conversely, WT1-TCR T cells showed only minimal cytotoxicity towards HLA-II^+^/WT1^+^ cell lines, similar to that of HLA-II^-^/WT1^-^ control (Fig. 4B), likely due to HLA mismatch (Supplementary Table S2).

We additionally performed cytotoxicity assays against primary AML or acute lymphoblastic leukemia (ALL) samples. In six out of nine samples screened, WT1-CAR induced significantly greater cytotoxicity as compared to the control (Fig. 4C). Similar to cell lines, recognition by WT1-CAR T correlated with the combined expression levels of WT1 and HLA-II on primary leukemic samples (Supplementary Fig. S7B).

### WT1-CAR T cells demonstrate antitumor activity *in vivo*

We next investigated the ability of WT1-CAR T cells to control tumor growth *in vivo*. HLA-II^+^ WT1^+^ PR9 cells were engrafted into immunodeficient mice. Two days post-engraftment, tumor-bearing mice were treated with WT1-CAR T cells or control mesothelin (MSLN)-CAR T cells (Fig. 5A). Consistent with our *in vitro* observations, WT1-CAR T cells showed a strong antitumor response, resulting in a slower progressed bioluminescence imaging signal (Fig. 5B and Supplementary Fig. S8) and better overall survival (Fig. 5C). Additionally, while MSLN-CAR treated mice exhibited circulating PR9 cells in peripheral blood, no PR9 cells were detected in mice treated with WT1-CAR T cells (Fig. 5D). Whereas MSLN-CAR T persistence fell markedly after 14 days post-treatment, WT1-CAR T cells could be detected at 28 days post-treatment (Fig. 5E).

**Figure 5:**
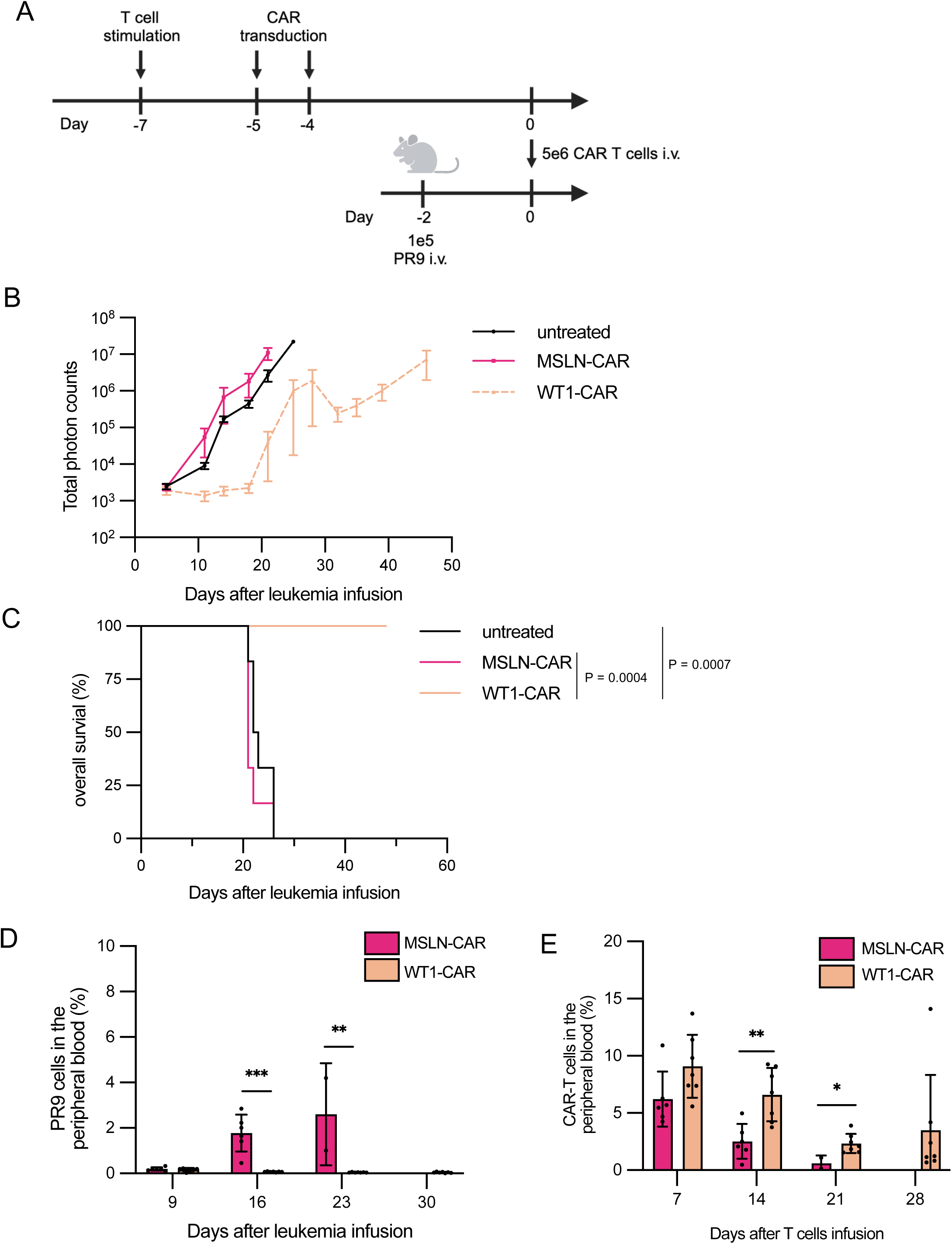
WT1-CAR demonstrates anti-tumor activity *in vivo*. (A) Experimental design showing NSG mice intravenously infused with 1 x 10^5^ luciferase-expressing PR9 cells (day -2) and adoptively transferred with 5 x 10^6^ CAR-T cells (day 0) or remained untreated. (B) Kaplan-Meier curve of the overall survival of mice in each treatment cohort. n = 6 mice per group. P values calculated by the log-rank test. (C) Tumor burden of mice with indicated treatment was analyzed by *in vivo* bioluminescent imaging of luciferase activity and total photon counts. n = 6 mice per group. (D) Persistence of PR9 cells in peripheral blood was analyzed by flow cytometry at the indicated time points. **P<0.01, *** P<0.001 by independent t-test for samples on each day. There was not a viable MSLN-CAR treated mouse available on day 30 for PR9 cell persistence analysis. (E) Persistence of CAR T cells in peripheral blood was analyzed by flow cytometry at the indicated time points. *P<0.05, **P<0.01 by independent t-test for samples on each day. There was not a viable MSLN-CAR treated mouse available on day 28 for CAR T cell persistence analysis.

## Discussion

A recent study identified a peptide-centric CAR using a phage-display counter panning strategy capable of recognizing the PHOX2B peptide cross-presented by HLA-A*23:01 and A*24:02 (31). While promising, this approach is limited by the biophysical constraints of the pHLA-I complex, which can only accommodate short peptides, reducing the availability of solvent-exposed residues. TCR mimetics developed by other groups make contact with both HLA-I and peptide simultaneously (3,32). Conversely, the open-ended binding groove of HLA-II molecules, which can accommodate longer peptides resulting in solvent accessible peptide-flanking residues (PFRs) that reside outside the HLA-II binding cleft (33), presents a target opportunity. By taking advantage of this property, we developed a WT1-CAR capable of recognizing WT1 peptide promiscuously presented across various HLA-II genotypes. As shown in our analyses, while a WT1-specific TCR could recognize WT1 presented by related HLA-DP alleles, our newly developed WT1-CAR was capable of recognizing WT1 across multiple HLA-DP, -DR, -DQ alleles. The WT1-CAR is derived from an antibody raised against the WT1 peptide alone, rather than the pHLA complex, resulting in cytotoxicity against various HLA-DP, -DR, and -DQ WT1-positive target cells. The ability to target WT1 presented by a wide range of HLA-II was necessitated by the unique targeting of the N-terminal PFR alone, but not portions in complex with the HLA molecules. Our analysis delineated that the N-terminal C330 and N331 are uniquely required for WT1-CAR recognition, while these residues did not appear to be critical for TCR interactions. Notably, as the 5H2 antibody was generated against the peptide in the absence of HLA complex, this finding suggests that the primary function of HLA-II in this context is to present an otherwise intracellular epitope on the cell surface, rather than contributing directly to CAR binding.

The broad presentation of WT1 peptide across multiple HLA-II alleles can be attributed to the conserved binding motifs characteristic within the peptide sequence. The primary HLA-II anchor pocket, P1 pocket, has been shown to preferentially accommodate large aromatic residues, and the consecutive presence of Y334 and F335 likely facilitates conserved and shifted binding registers across different HLA-II molecules (34). Our findings further identify F335 and L340 as primary anchor residues for HLA-DP4 binding, aligning with previous reports that describe a uniform anchor spacing pattern within the HLA-DP alleles. Specifically, the predominant anchor residues for HLA-DP alleles are positioned at P1 and P6, which are essential for stable peptide binding (35). Notably, WT1-CAR T cells failed to recognize the WT1 peptide presented by HLA-DR1301. HLA-DR1301 has been reported to exhibit a distinct binding preference, favoring smaller hydrophobic residues such as valine at the P1 pocket, rather than the bulkier tyrosine or phenylalanine residues (36). This distinct binding specificity suggests that HLA-DR1301 may not accommodate WT1_330–348_ in a register compatible with CAR engagement, thereby impairing 5H2 recognition. In addition, M342 was identified as a key residue for HLA-DR1501 and -DQ9.2 binding, and it likely contributes to binding stability and sits within the P9 pocket. Interestingly, for HLA-DR1501, R333 and Y334 were also identified to be important for peptide loading, despite being solvent-exposed and located outside of the binding groove in the AlphaFold3-modeled structure. Analysis of the modeled complex revealed the formation of multiple hydrogen bonds within the N-terminal PFR, including interactions involving R333 and Y334.

These stabilizing interactions within the N-terminal PFRs are consistent with previous studies, which highlight that stable structures formed within PFRs have been shown to facilitate peptide-MHC binding affinity (37,38). Our findings suggest that although R333 and Y334 are not classical anchor residues, their contribution to PFR-mediated stabilization is critical for optimal peptide presentation by HLA-DR1501. Overall, our analysis reveals a largely conserved peptide-binding motif for the WT1 peptide across representative alleles from different HLA-II families. In particular, residues _334_YF_335_ and _340_LQM_342_ define a critical “hot zone” for peptide anchoring and structural stability. These results underscore the importance not only of primary anchor residues but also of adjacent, solvent-facing residues in shaping peptide conformation, influencing both peptide–MHC affinity and presentation (39,40).

To the best of our knowledge, this paper presents the first attempt to target an HLA-II-presented cancer antigen in an HLA-agnostic manner. Existing TCR-T cell therapies necessitate precise matching of HLA genotypes for patient eligibility, thereby limiting the range of individuals who can benefit from these treatments. Taking advantage of the naturally limited expression of HLA-II molecules in healthy tissues throughout the body, combined with an increase of HLA-II expression in tumors, our therapy was able to target and eliminate cancer cells while leaving relevant healthy cells unharmed, such as primary human HLA-II^+^ WT1^+^ CD34^+^ HPCs. This finding suggests that WT1 may be differentially processed in normal cells compared to tumor cells. Further preclinical studies will be necessary to evaluate off-tumor toxicity towards other reported WT1^+^ cells, such as podocytes and mesothelial cells, especially under a cancer or treatment-induced hormone and/or cytokine boost that may upregulate HLA-II molecules. In addition, WT1-CAR had non-detectable responses to highly homologous peptides, which, taken together with the HLA-agnosity, suggests the WT1-CAR we have generated holds the advantage of less “off-target” cross-reactivity, given TCR cross-reactivity can be accounted for by the structure mimicry induced by peptide-homology or MHC-focused binding (5,41,42).

Our peptide-specific HLA-II-restricted-yet-independent therapeutic strategy is poised to be applicable to other intracellular antigens presented by HLA-II. Notably, the germline cancer antigens NY-ESO-1 and MAGE-A3 are prime candidates due to the promiscuous presentation of their antigen epitopes across various HLA-II alleles (11,43,44). Further, the expression of these antigens in normal tissues is restricted mainly to germline cells and trophoblastic cells devoid of HLA-II expression, reducing the possibility of treatment-induced on-target, off-tumor toxicity (45).

In summary, we have developed an innovative approach to CAR-T therapy that can target peptides promiscuously presented across a variety of HLA-II alleles. The novel CAR construct has a peptide-centric, HLA-II-agnostic nature, which circumvents conventional HLA restrictions. This enhancement may expand therapeutic potential, making it accessible to a broader and more genetically diverse group of patients.

## Methods

### Cells lines

Peripheral blood samples were obtained from healthy donors following Institutional Review Board approval. Written and informed consent was collected from all donors. K562, HL60, Daudi, Ramos, U937, and PR9 cell lines were cultured in RPMI1640 (Gibco) supplemented with 10% fetal bovine serum (FBS, Gibco) and 0.1% gentamycin. KG1a and T2 cells were cultured in IMDM supplemented with 20% FBS and 0.1% gentamycin. OCI-AML5 were cultured in alpha-MEM (Gibco) supplemented with 20% FBS, 10 ng/ml human GM-CSF and 0.1% gentamycin. Daudi, Ramos, HL60, T2, and K562 were obtained from the American Type Culture Collection (ATCC). KG1a, OCI-AML5, U937, PR9, and primary AML/ALL samples were provided by Dr. Mark Minden through the Leukemia Tissue Bank at Princess Margaret Cancer Centre, University Health Network, Toronto, Canada. Primary human CD34^+^ hematopoietic cells purified from cord blood samples (STEMCELL Technologies).

### Genes

WT1-TCR⍺β genes were fused by a furin cleavage site, an SGSG spacer sequence, and an F2A sequence(furin-SGSG-F2A) (21,46), followed by furin-SGSG-F2A linked to a truncated version of nerve growth factor receptor (ΔNGFR). Immunoglobulin genes of 5H2 mAb were cloned via 5’-rapid amplification of cDNA ends (5’-RACE) PCR as previously described(47). The single-chain variable fragment (scFv) consists of the variable regions of heavy chain (V_H_) and light chain (V_L_) derived from 5H2 and a Whitlow linker (48). The scFv (FMC63-derived for CD19 CAR, SS1-derived for MSLN CAR) was linked to human CD28 transmembrane and cytoplasmic domains, and the cytoplasmic domain of human CD3ζ (49,50). The CAR constructs and full-length WT1 cDNA were individually N-terminally linked to ΔNGFR via a furin-SGSG-P2A (51).

All cDNAs were cloned into the pMX retroviral vector, and all transduced cells were generated using retrovirus produced by 293GPG or PG13 cells as previously published (52,53). HLA-II expressing K562 cells were generated using retrovirus transduction (54), and individually transduced with the following HLA-II genes: *DPA1*01:03/DPB1*01:01 (DP1), DPA1*01:03/DPB1*02:01 (DP2), DPA1*01:03/DPB1*03:01 (DP3), DPA1*01:03/DPB1*04:01 (DP4-0401), DPA1*02:01/DPB1*04:02 (DP4-0402), DPA1*02:01/DPB1*05:01 (DP5), DPA1*02:01/DPB1*13:01 (DP13), DPA1*02:06/DPB1*104:01 (DP104), DQA1*02:01/DQB1*02:02 (DQ2.2), DQA1*01:02/DQB1*05:02 (DQ5.2), DQA1*01:04/DQB1*05:03 (DQ5.3), DQA1*02:01/DQB1*03:03 (DQ9.2), DRA1*0101/DRB1*01:01 (DR1), DRA1*0101/DRB1*04:01(DR4), DRA1*0101/DRB1*07:01(DR7), DRA1*0101/DRB1*13:01(DR13), DRA1*0101/DRB1*14:54(DR14), DRA1*0101/DRB1*15:01(DR1501), DRA1*0101/DRB1*15:02(DR1502), DRA1*0101/DRB1*16:01(DR16), DRA1*0101/DRB5*02:02 (DR51b), DRA1*0101/DRB3*02:02 (DR52b)*. HLA-II levels were confirmed by flow cytometry. HLA-II^+^ K562 cells were transduced with WT1/ΔNGFR to ectopically overexpress WT1. Cells transduced with constructs containing ΔNGFR were further purified using anti-NGFR beads (Miltenyi Biotec). eGFP-luciferase-expressing cell lines were generated by retroviral transduction of a pMX-EGFP–firefly luciferase construct. HLA-II genotyping was conducted through Scisco Genetics Inc.

### Transduction of T cells

CD3^+^ T cells were purified using the Pan T Cell Isolation Kit (Miltenyi Biotec) and stimulated at an E:T = 5:1 for 48 h with aAPC/mOKT3 that had been irradiated with 200 Gy for 90 mins (54,55). PG13 cell lines stably transduced with TCR- or CAR-expressing plasmid were used for T cell transductions. PG13 virus was added to Retronectin-coated plates (Takara Bio) and centrifuged at 2,000 x*g* for 2 h at 32 °C for two consecutive days. T cells were suspended in medium containing 100 IU/ml IL-2 and 10 ng/ml IL-15 and added to the plates. The culture medium was replenished every 2–3 days.

### ELISA assays

ELISA plates were coated with anti-His antibodies and incubated at 4°C overnight. The next day, the plates were washed and coated with DP4/CLIP or DP4/WT1_329–348_ monomers, or CLIP (_97_LPKPPKPVSKMRMATPLLMQALPM_120_) or WT1_330-348_ (_330_CNKRYFKLSHLQMHSRKHT_348_) and incubated at 4 °C overnight. The monomers were procured from the National Institutes of Health (NIH) Tetramer Core Facility. The next day, the plates were washed extensively before adding WT1-specific or control mAb. After incubation at room temperature for 2 h, plates were washed extensively and incubated with ALP-anti-mouse IgG/IgM at room temperature for 30 mins. Finally, the plates were washed and incubated with p-nitrophenyl phosphate (PNPP) substrate (Pierce, Rockford, IL) at room temperature. The reaction was terminated by adding 1 M NaOH. The optical density (OD) (405 nm) was read (Spectramax 190 Microplate Reader; Molecular Devices, Sunnyvale, CA).

### Biolayer interferometry (BLI)

BLI experiments were performed using an Octet Red96 instrument with streptavidin (SA)-coated biosensor tips (Sartorius). All experiments were performed at 25 °C with black flat-bottom 96-well microtiter plates (Greiner Bio-One GmbH, Germany). SA tips were hydrated in HPS-EP^+^ buffer (10 mM HEPES, pH 7.4, 150 mM NaCl, 3 mM EDTA, 0.005% Tween-20; Cytiva, Sweden AB) for 10 mins. Following sensor check measurements in HPS-EP^+^ buffer, tips were dipped into 200 µL of biotinylated reagents diluted in HPS-EP^+^ buffer until approximately 2.0 nm response was achieved. The biotinylated reagents used include 50 nM of N-linked biotinylated WT1_330-348_ peptide (Genscript, Piscataway, NJ), biotinylated *DPB1*04:01/DPA1*01:03*-linked human CLIP_87-101_ (PVSKMRMATPLLMQA) (NIH) and biotinylated *DPB1*04:01/DPA1*01:03*-linked WT1_329-348_ (GCNKRYFKLSHLQMHSRKHT) (NIH). Peptide-loaded SA tips were then quenched in a 100 nM solution of free biotin (Sigma-Aldrich) for 120 s, followed by baseline measurements in the buffer to remove excess peptide for an additional 120 s. The association step with 5H2 antibody at various concentrations was performed for 600 s, followed by dissociation for 600 s in buffer only. Data were analyzed using Octet Data Analysis software and steady state affinity measurements at all concentrations were globally fit using a Langmuir (1:1) model.

### Flow cytometry

For surface staining, cells were pelleted and resuspended in a master mix diluted in PBS/2% FBS. Cells were incubated at 4°C for 15 mins and washed before analysis. The following antibodies were used for surface staining: CD3 (UCHT1, BioLegend), CD4 (RPA-T4, BioLegend), CD8 (RPA-T8, BioLegend), pan HLA class II (Tü39, BioLegend), HLA-DP (B7/21, Leinco Technologies), HLA-DR (L243, BioLegend), HLA-DQ (Tü169, BioLegend), NGFR (ME20.4, BioLegend), PD-1 (EH12.2H7, BioLegend), Tim-3 (F38-2E2, BioLegend), LAG-3 (7H2C65, BioLegend), and mouse isotype controls (BioLegend). For intracellular WT1 expression detection, target cells were fixed and permeabilized using the Cytofix/Cytoperm Kit (BD Biosciences) and stained with mouse anti-WT1 mAb (Novus Biologicals), followed by BV421-anti-mouse IgG (Poly4053, BioLegend). Surface expression of CAR was confirmed by biotinylated protein-L (GenScript) (56).

### ELISpot assays

IFN-γ ELISpot was performed as previously described (52). Briefly, T cells and 2 x 10^4^ target cells were co-cultured for 20-24 h at 37 °C on polyvinylidene difluoride (PVDF) plates (Millipore) coated with IFN-γ capture monoclonal antibody (1D1K, MABTECH). Plates were washed and incubated with biotin-conjugated IFN-γ detection monoclonal antibody (7-B6-1, Mabtech). HRP-conjugated streptavidin (DAKO) was added for substrate development. The peptides used included WT1_328-348_ (_328_PGCNKRYFKLSHLQMHSRKHT_348_), alanine or deletion mutants thereof, and cross-reactive peptide candidates (Supplementary Table S2). All peptides were purchased from GenScript.

### *in vitro* cytotoxicity assays

K562 cells were labeled with 5 μM fluorescent Vybrant™ DiO in PBS (Thermo Fisher Scientific) at 1e6/mL for 15 mins at 37 °C, washed and added to 96-well plates in 100 μL culture medium. TCR or CAR T cells were added at the indicated E:T ratio. After 6 h or 18 h co-culture, cell mixtures were stained with 3 μM TO-PRO-3 (Thermo Fisher Scientific) for 30 mins at room temperature before flow cytometric assays. Live target cells after co-culture were defined as % of DiO^+^ TO-PRO-3^+^ cells in the culture. Cytotoxicity was defined as Cytotoxicity = (% live target cells without effector cells -% live target cells with effector cells)/ % live target cells without effector cells.

### Competitive binding assays

T2 cells were washed in serum-free IMDM (Gibco) and incubated at 37°C for 3 h with 1 µM biotinylated-CLIP peptide, either alone or in the presence of competitive peptides at the indicated concentrations. Cells were resuspended at 5 × 10⁵ cells/50 µL in IMDM medium. Following incubation, cells were washed with PBS/2% FBS and subsequently stained with 10 µg/mL streptavidin-PE (Invitrogen) at room temperature for 30 mins. Flow cytometric analysis was then performed. Dose-inhibition fitting was applied to the resulting curve. LogIC50 and R^2^ of the fitted curve were calculated (Prism 10.4.1). Mutants were deemed “poor binders” when LogIC50 >1 or R^2^ <0.9.

The percentage of inhibition was calculated based on median fluorescence intensity (MFI) using the formula: % Inhibition = [1-(MFI with CLIP peptide only/MFI with competitive peptide)] x 100.

### Prediction algorithms

HLA binding modeling was created using AlphaFold3 (27). WT1_330-348_ sequence (CNKRYFKLSHLQMHSRKHT) and HLA-II ⍺β chain protein sequences (IPD-IMGT/HLA database) were used as inputs. The top-ranked model, confirmed with iPTM score >0.7, was visualized using PyMOL. Hydrogen bonds were identified by analyzing polar contacts between chains and within individual chains, using a distance cutoff of 3.6 Å. HLA surface macromolecular electrostatics was calculated using the Adaptive Poisson-Boltzmann Solve (APBS) algorithm.

Red: negative charge; Blue: positive charge. For BLAST searches, WT1_330-348_ sequence was used as input for NCBI protein BLASTp for Homo sapiens protein entries in the RefSeq: NCBI Reference Sequence Database. Results were filtered by >50% identity and >50% query coverage and sorted by low E-values and high max score before top candidates were selected.

### xCELLigence assays

Long-term, continuous T cell-mediated cytotoxicity assays were conducted on the xCELLigence Real-Time Cell Analyzer (RTCA)-Multiple Plate instrument (Agilent Technologies). The impedance-based analysis was conducted according to the manufacturer’s instructions. Briefly, 96-well plates (E-Plate^®^View 96, Agilent Technologies) were incubated at 4 °C overnight with 4 µg/mL tethering reagent (anti-CD71 tethering kit). Plates were washed, applied with culture medium, and allowed to equilibrate in the instrument at 37 °C for 60 mins before initiating background measurements. Target cells were then added to each well at indicated E:T ratios in a total of 200 µL culture medium. T cells were added on the following day. The plate impedance was measured every 15 mins after experiment initiation. Cell index values were calculated by RTCA software based on electrical impedance and converted to percent cytotoxicity: % Cytotoxicity = [(cell index without T cell - cell index with T cells)/cell index without T cells] x 100. For fluorescence signal-based analysis, the eGFP-transduced target cells were combined with effector cells at the indicated E:T ratio at time zero. Total fluorescence intensity (TFI) was measured every 90 mins and normalized to the first measured timepoint. %Cytotoxicity= [(TFIwithout T cells −TFIwith T cells)/TFIwithout T cells] × 100.

### Time-of-flight mass cytometry (CyTOF)

Long-term co-cultures were set up at an E:T ratio of 1:3, and samples were submitted for CyTOF analysis on the first, third and fifth day after the initiation of co-culture. CyTOF staining was performed as previously described, using antibodies directly conjugated to metal tags (57). Briefly, samples were washed with PBS and incubated with 1 µM Cell-ID Intercalator 103Rh in HBSS for live/dead discrimination for 15 mins at 37 °C. Cells were fixed with eBioscience Foxp3 Fixation/Permeabilization reagent for 15 mins at room temperature, washed with 1X Barcoding Permeabilization buffer and incubated with a unique palladium barcode for 30 mins at room temperature. Following several washes, barcoded samples were pooled into a single tube and FcR blocked for 10 mins prior to incubation with a cocktail of surface metal-tagged antibodies prepared in PBS/2% FBS. After 30 mins at 4 °C, the pooled sample was washed and incubated for 30 mins at 4 °C with a cocktail of intracellular metal-tagged antibodies prepared in 1x Permeabilization buffer (eBioscience). DNA staining with 125 μM Cell-ID Intercalator Ir was performed for 1 h at 4 °C to distinguish whole cells from debris. Samples were stored in PBS/1.6% paraformaldehyde until the time of acquisition. Samples were acquired on a Helios mass cytometer at the PMCC Centre for Integrative Immune Analysis Facility. The CyTOF panel used is as follows:

| Channel | Panel | Channel | Panel | Channel | Panel |
| --- | --- | --- | --- | --- | --- |
| 89 Y | CD45 | 154 Sm | CD69 | 170 Er | CTLA-4 |
| 103 Rh | viability | 155 Gd | CD45RO | 171 Yb | Helios |
| 141 Pr | CD45RA | 158 Gd | CD27 | 172 Yb | CD127 |
| 142 Nd | HLA-DR | 159 Tb | CCR7 | 173 Yb | CD56 |
| 143 Nd | ICOS | 160 Gd | ISG15 | 174 Yb | TCF-1 |
| 145 Nd | CXCR3 | 162 Dy | CD28 | 175 Lu | PD-L1 |
| 146 Nd | CD8a | 163 Dy | 4-1BB | 176 Yb | LAG3 |
| 147 Sm | CD4 | 164 Dy | Granzyme B | 191 Ir | DNA |
| 149 Sm | FOXP3 | 165 Ho | PD-1 | 193 Ir | DNA |
| 150 Nd | CD103 | 166 Er | TIM3 | 209 Bi | TIGIT |
| 151 Eu | CD39 | 167 Er | CD95 | 114 Cd | BST2 |
| 153 Eu | CD3 | 168 Er | Perforin | 116 Cd | Ki67 |

Bead-normalized FCS files were pre-processed using FlowJo v10.8.1 software. Samples were debarcoded and exported as individual FCS files. A series of sequential gates was applied to exclude beads, doublets, ion fusions and dead cells, followed by gating on CD45^+^CD3^+^ events. CD45^+^CD3^+^ single-cell intensities were then exported as CSV files and imported into R for downstream analysis. Data were transformed using hyperbolic arcsine with a custom scaling factor for each channel. Dimensionality reduction was performed using the “umap-learn” method from the package *umap* (v0.2.7.0) in R (v4.0.0), and unsupervised clustering was performed using the fast-Phenograph algorithm using the *Fast-PG* package (v0.0.6) in R (v3.6.2). All subsequent CyTOF analyses were performed in R (v4.2.1). Data visualizations were generated using functions available in *ggplot2* (v3.4.0), *pheatmap* (v1.0.12) *ggh4x* (v0.2.3), and *ggpubr* (v0.4.0) packages.

### Mouse experiments

Six- to eight-week-old male NOD-scid-IL2Rg^null^ (NSG) mice were purchased from The Jackson Laboratory and bred and housed at the Princess Margaret Cancer Center Animal Facility (ARC). Mice were first irradiated with 1.5 Gy using X-RAD 320 (Precision X-Ray) before the transplantation of 1 x 10^5^ eGFP-luciferase-expressing PR9 leukemia cells through intravenous (i.v.) injections. NSG mice received 5 x 10^6^ CAR-T cells (i.v.) 48 h post-leukemia engraftment. Mice were injected intraperitoneally with 200 µL of 15 mg/mL D-luciferin in PBS (Caymen Chemical) and imaged with Xenogen IVIS Spectrum (PerkinElmer) 20-30 mins post-injection. Mice were monitored at least four times per week and euthanized when they became moribund due to leukemia progression or reached endpoints outlined in institutional guidelines. The mice were randomly assigned to treatment groups in each experiment. No statistical methods were used to predetermine the sample size. The investigators were not blinded to allocation during experiments and outcome assessment.

### Statistical analyses

Statistical analyses were performed using R (for CyTOF data) and GraphPad Prism. For mouse experiments, survival was compared by the log-rank test. No statistical method was used to predetermine the sample sizes. The investigators were not blinded to allocation during experiments and outcome assessment. For CyTOF data, tests for differential abundance of immune cell clusters were performed using two-way ANOVA for multi-factor comparisons, Wilcoxon or the edgeR test from the *diffcyt* R package (v1.16.0) for comparison of two groups.

## Acknowledgments

We thank Hobin Seo, Ramy Gadalla, and the Princess Margaret Cancer Centre Flow Cytometry Facility for their technical support. N.H. was the Amgen Chair in Cancer Research and is the Tier 1 Canada Research Chair in Immunology to Immunotherapy. This work was supported by the Canadian Institutes of Health Research Project Grant PJT148586 (to N.H), the Ontario Institute for Cancer Research Clinical Investigator Award IA-039 (to N.H.), the Longo Family Cancer Foundation (to N.H.), the Ira Schneider Memorial Cancer Research Foundation (to N.H.), the Princess Margaret Cancer Foundation (to N.H.), the Longo Family Cancer Immunotherapy Fellowship (to Y.M.), and the Frederick Banting and Charles Best Canada Graduate Scholarship (to C.-H.W.). The experimental schematics were created with BioRender.com.

## Author Contributions

C.-H.W., E.Y.F.Z., T.O., B.X.W. and N.H. designed the research. C.-H.W., E.Y.F.Z., T.O., Y.O., F.I.,

S.F., G.M.B., D-H.H., X.W., B.D.B., K.S., Y.M., and D.L. performed the experiments. Y.K., M.O.B. and M.D.M. provided critical reagents. All the authors analyzed the results. C.-H.W., E.Y.F.Z., B.D.B., D.L., and N.H. designed the figures and drafted the manuscript. All the authors reviewed and contributed to the manuscript.

---

**Supplementary Figure 1:**
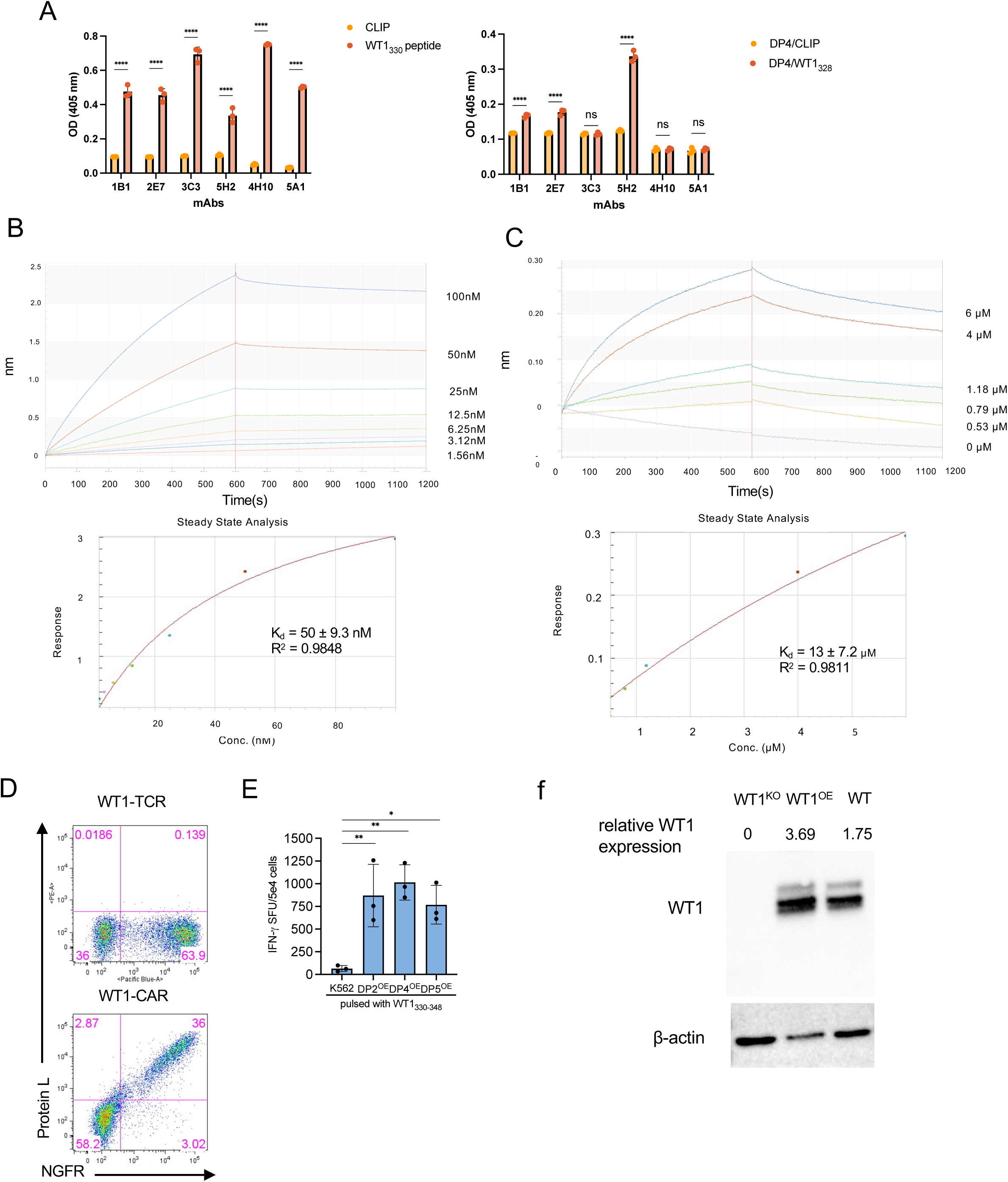
WT1-specific mAb binds to WT1 peptide presented by HLA-DP molecules. (A) Binding of WT1-specific mAbs to immobilized WT1 peptide or DP4/WT1 monomers was assessed by ELISA. Left: mAbs were added to the ELISA plate coated with WT1_330-348_ or CLIP (control). Right: mAbs were added to the ELISA plate coated with DP4/WT1330-348 monomers or DP4/CLIP monomers (control). The data shown represent the mean ± SD of experiments performed in triplicate. Results are representative of two independent experiments.ns, not significant, *p<0.05, **p<0.01, ***p<0.001, ****p<0.0001 by two-way ANOVA with Bonferroni tests. (B) Quantification of binding between WT1-specific mAb, 5H2, and WT1_328-348_. Interaction strength was measured by a biolayer interferometry (BLI) binding assay. (C) Quantification of binding between WT1-specific mAb, 5H2, and DP4/WT1_328-348_ monomers. Interaction strength was measured by a BLI binding assay. (D) Flow cytometry plot of surface CAR expressions (stained by protein L) versus tag (ΔNGFR) expressions. (E) IFN-γ secretion by WT1-TCR T cells against K562 cells expressing the indicated HLA-II molecules pulsed with WT1_328-348_ at an E:T ratio of 5:1 for 20-24 h measured by ELISpot. n = 3 donors. (F) Western blot analysis of WT1 expressions in K562/WT1^KO^, K562/WT1^OE^, and wild-type K562 (wt).

**Supplementary Figure 2:**
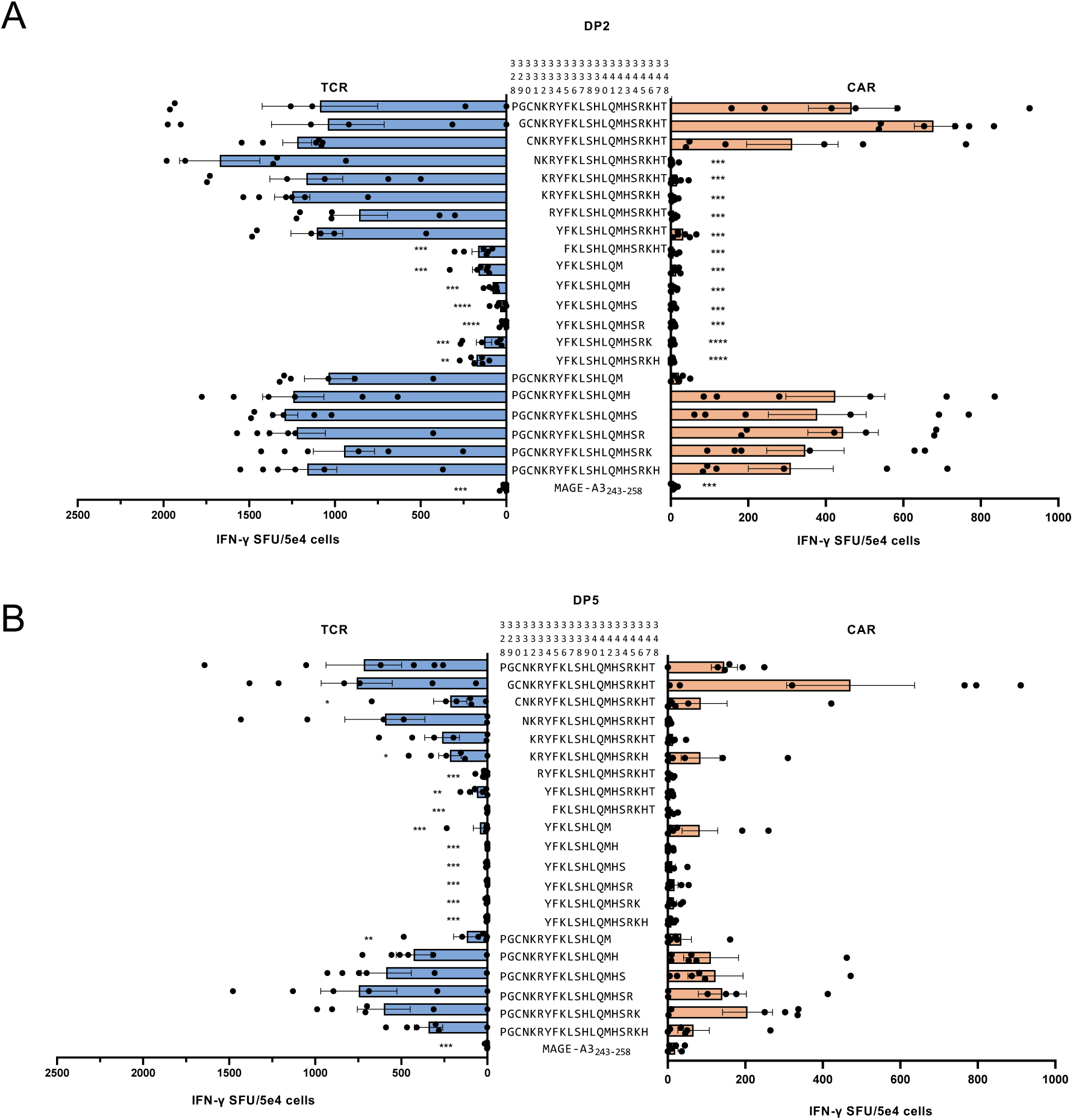
WT1-TCR and CAR have distinct modes of recognition of WT1 peptide presented with a homologous binding motif across HLA-DP. WT1-TCR or CAR T cells were stimulated with (A) T2/DP2 or (B) T2/DP5 pulsed with different deletion peptides, for 20-24 h. MAGE-A3_243-258_ was used as a negative control. IFN-γ secretion was measured by ELISpot analysis. n = 6 donors. *P<0.05, **P<0.01, ***P<0.001, ****P<0.0001 by one-way ANOVA multiple comparison test with Bonferroni tests.

**Supplementary Figure 3:**
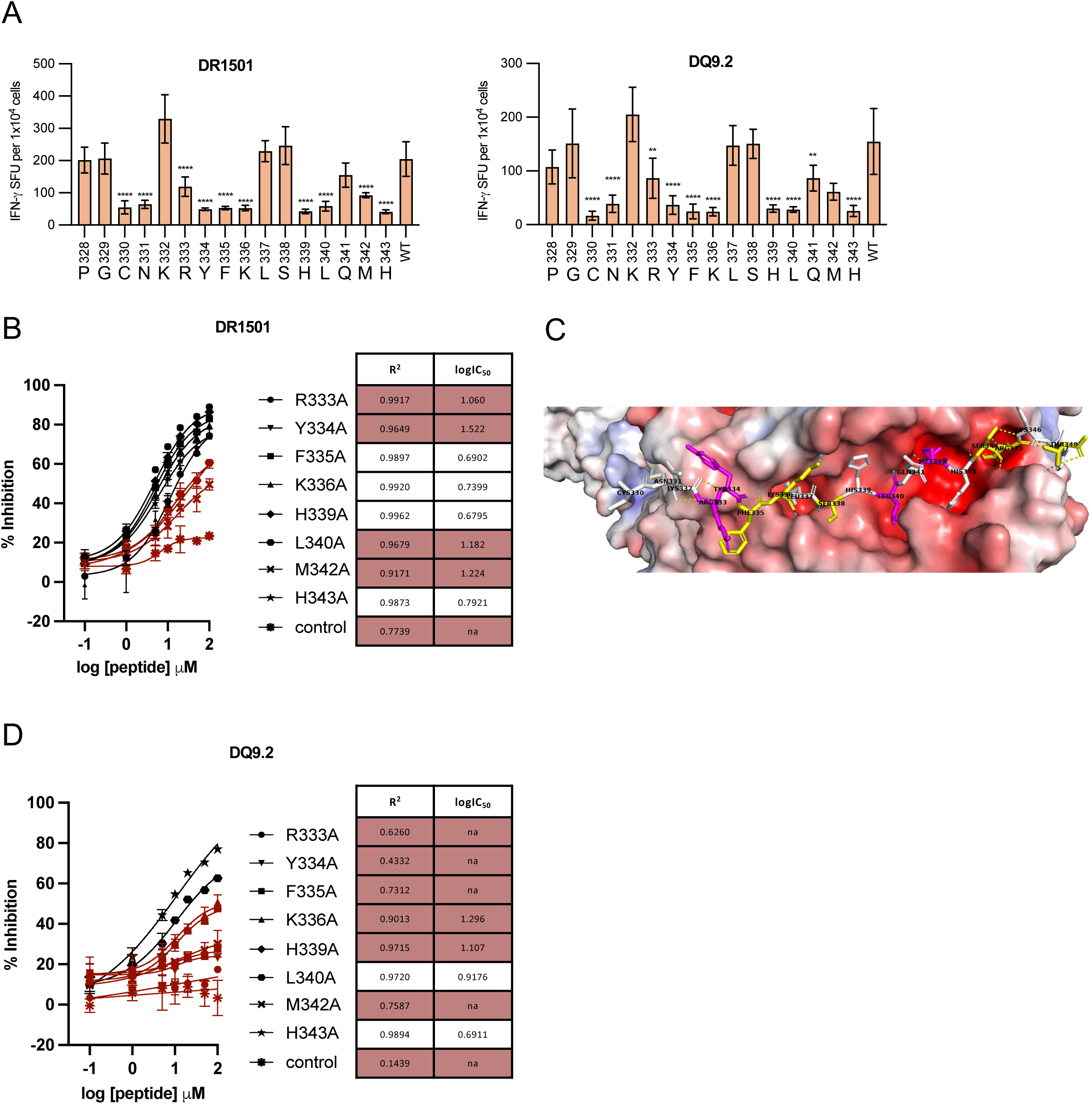
WT1 is presented with a highly conserved motif across HLA-II families. (A) WT1-CAR T cells were stimulated with T2/DR1501 or T2/ DQ9.2 pulsed with WT1_328-348_ (WT) or WT1_328-348_ substituted with alanine at indicated positions for 20-24 h. IFN-γ secretion was measured by ELISpot. n = 3 donors. (B) Competitive binding assay with T2 cells expressing HLA-DR1501, pulsed with graded concentration of WT1_328-348_ alanine mutant at specified positions in the presence of 1 µM biotinylated-CLIP peptide. Poor binders (Methods) are marked in red. Data showing the mean ± SD from 3 independent studies. (C) Structural modelling of WT1_330-348_ presented in HLA-DR1501 by AlphaFold3 showing HLA surface electrostatic charge (red: negative, blue: positive), peptide-HLA H-bonds (light blue dash), intra-peptide H-bonds (yellow dash), anchors identified in 2B (magenta), residues forming H-bonds (yellow).

**Supplementary Figure 4:**
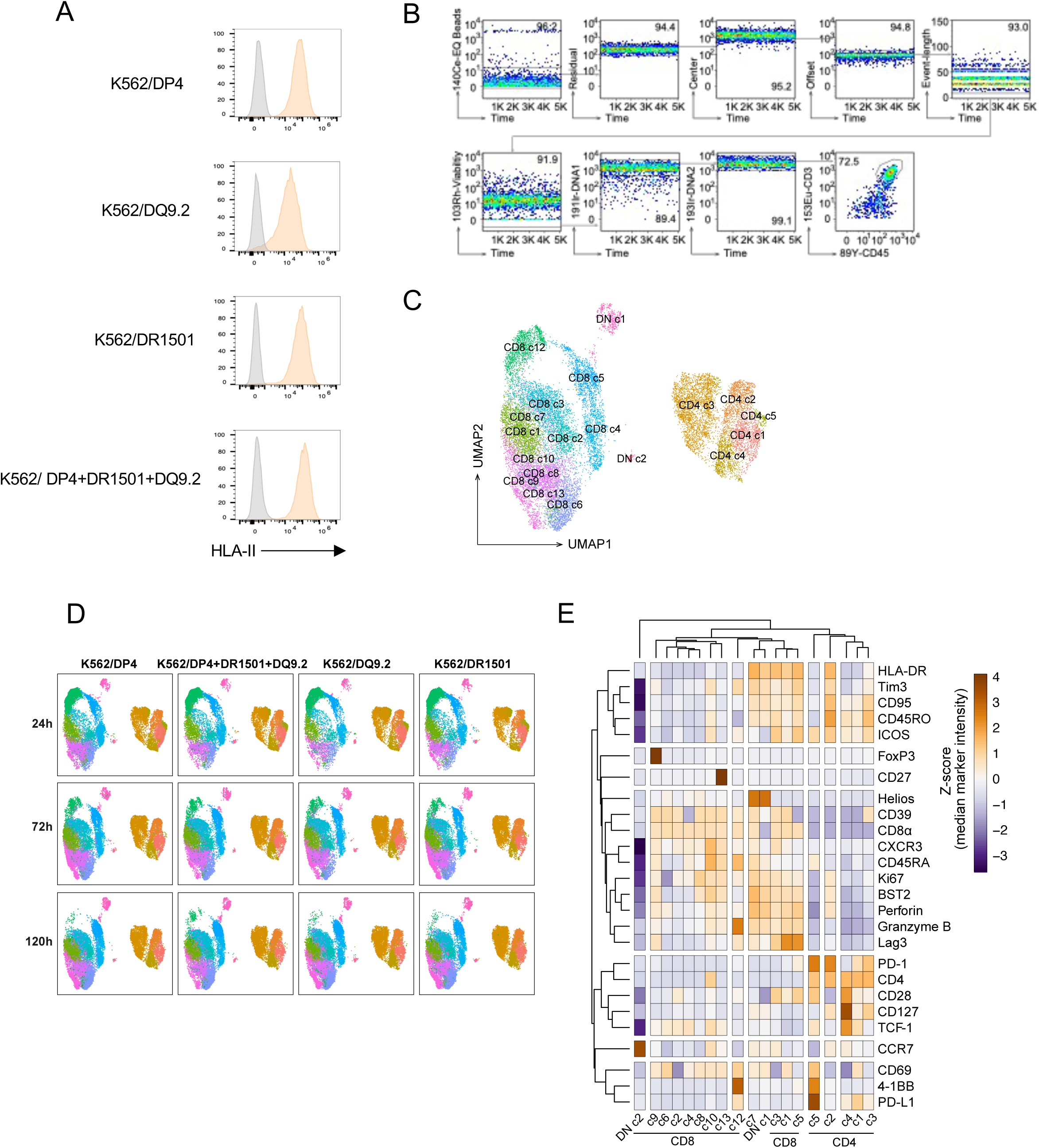
WT1-CAR T cells targeting cells expressing multiple HLA-II molecules. (A) HLA-II surface expression on K562 cells transduced with indicated HLA-II molecules analyzed by flow cytometric assays staining with anti-HLA-II mAb. (B) Flow cytometric gating for CD45^+^CD3^+^ CAR T cell population before CyTOF analysis for Figure 2(B). (C) UMAP plots for all profiles from CyTOF, colored according to the identified cell types. (D) UMAP plots of all identified populations in CAR T cells stimulated with different target cells across timepoints. (E) Heatmap of the median marker intensities of all the lineage and phenotype markers across the 19 cell populations identified in (B) (excluding CD8 c11, identified to be T cell-K562 cell doublets population).

**Supplementary Figure 5:**
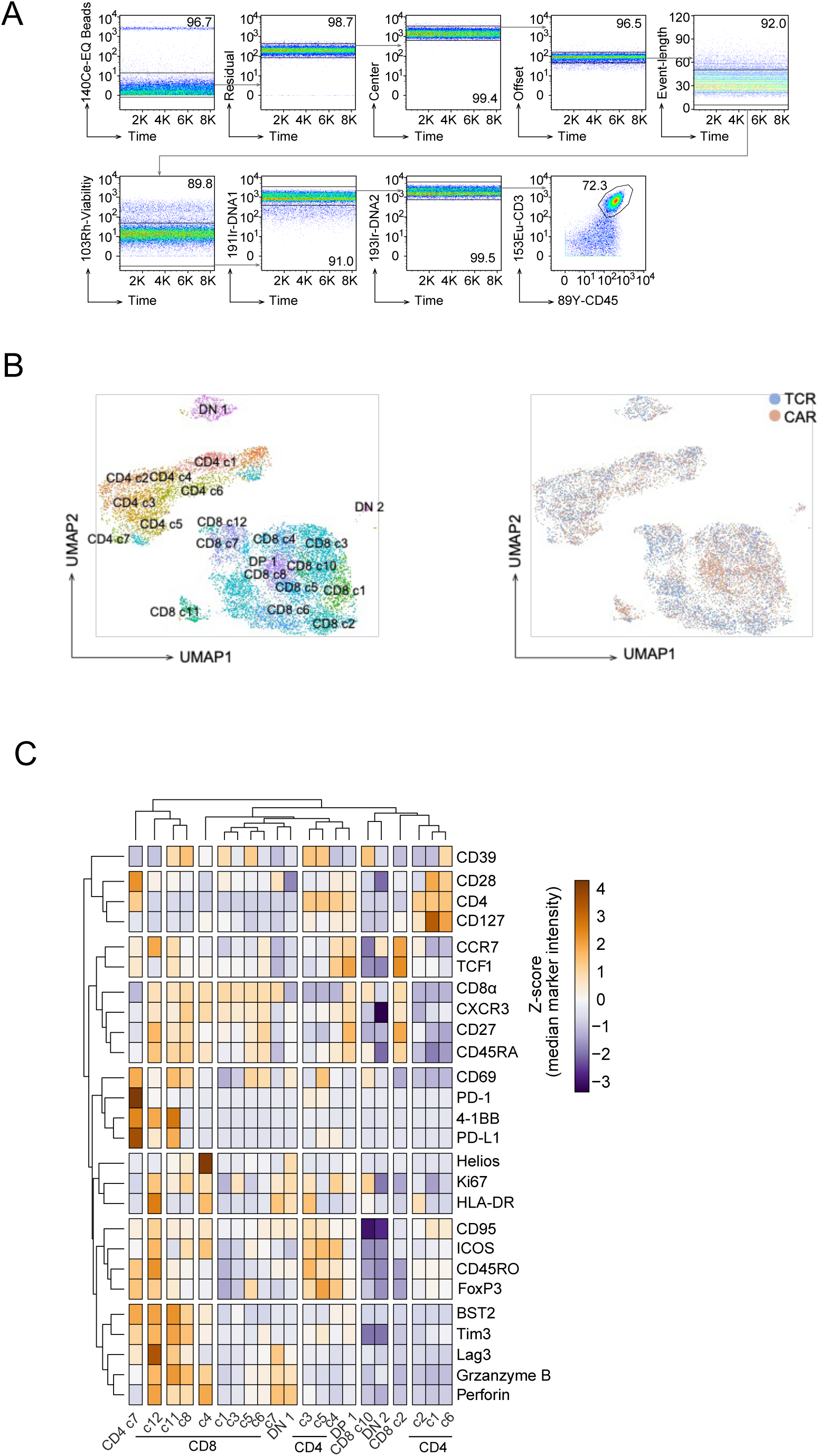
WT1-CAR and -TCR T cells show distinctive phenotypic changes upon stimulation by target cells with multiple HLA-II molecules. (A) Flow cytometric gating for CD45^+^CD3^+^ T cell population before CyTOF analysis for Fig. 2 (D) and (E). (B) (Left) UMAP plots for all profiles from CyTOF, colored according to the identified cell types. (Right) UMAP plots for all profiles showing origin of cells (WT1-TCR or WT1-CAR). (C) Heatmap of the median marker intensities of all the lineage and phenotype markers across the 21 cell populations identified in (B) (excluding CD8 c9, DN 3, identified to be T cell-K562 cell doublets populations).

**Supplementary Figure 6:**
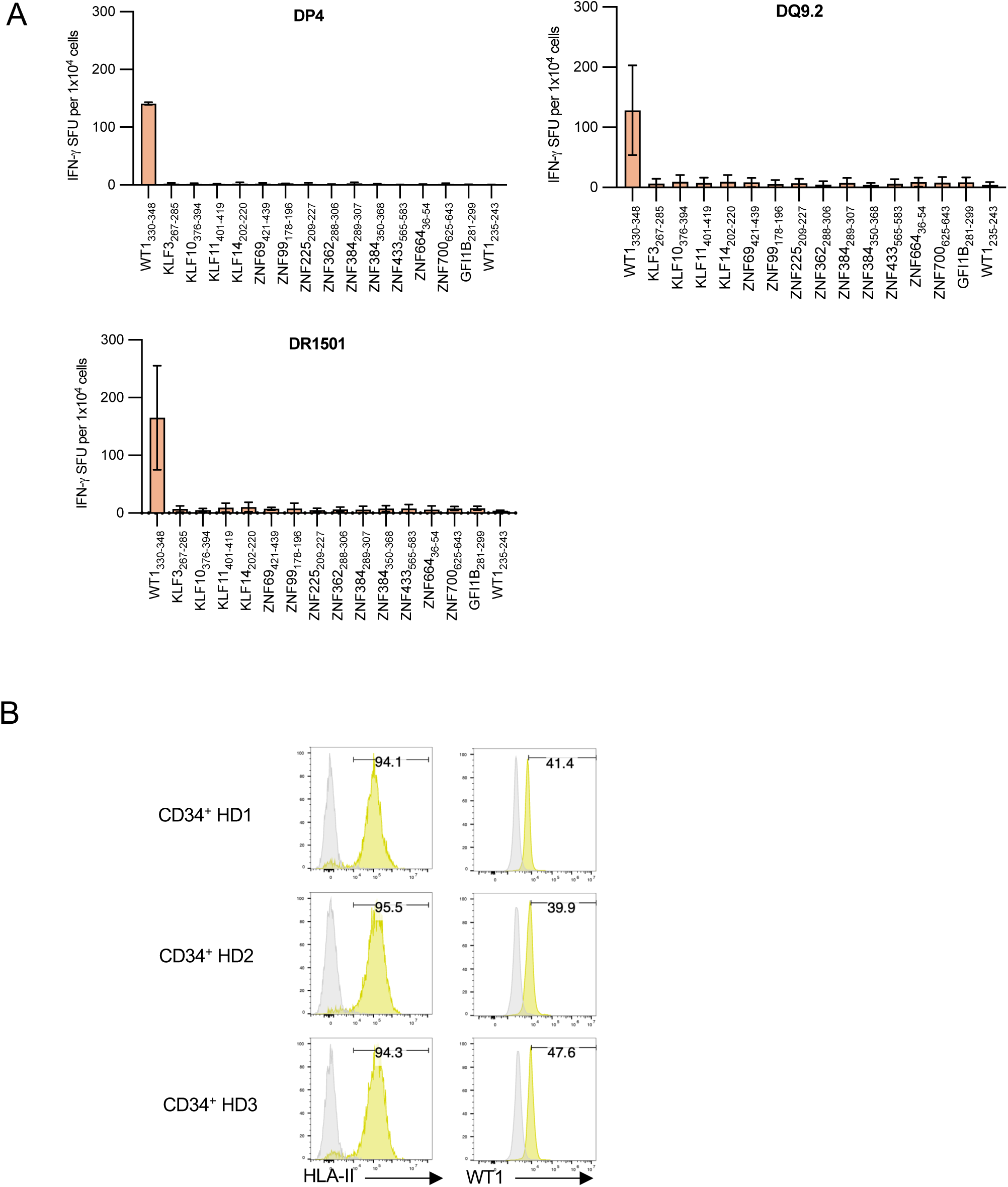
WT1 CAR T cells show minimal cross-reactivity and “on-target, off-tumor” toxicities. (A) WT1-CAR T cells were stimulated with T2 cells expressing the indicated HLA-II molecules pulsed with WT1_330-348_ or WT1_235-243_ (control) or cross-reactive peptide candidates for 20-24 h (Supplementary Table S3); IFN-γ secretion was measured by ELISpot analysis. The data shown represent the mean ± SD of experiments performed in triplicate. Results are representative of two independent experiments. (B) Surface HLA-II and intracellular WT1 expressions in primary human CD34^+^ hematopoietic samples from three different healthy donors were measured by flow cytometric analysis after staining with isotype control, anti-HLA-II mAb or anti-WT1 mAb.

**Supplementary Figure 7:**
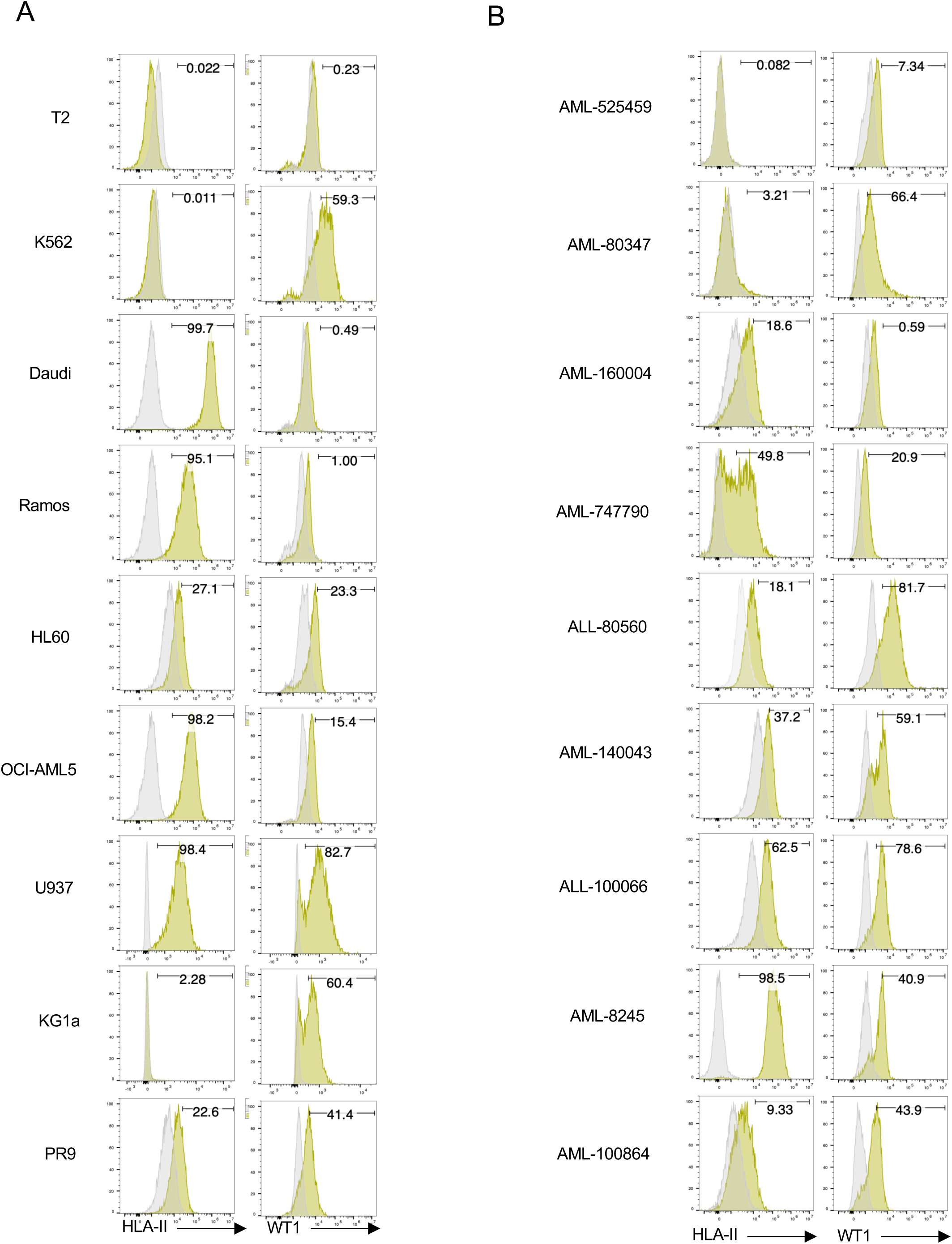
HLA-II and WT1 expression of a panel of ALL or AML samples and cell lines. Expressions of surface HLA-II and intracellular WT1 proteins of (A) cell lines and (B) primary ALL or AML samples were measured by flow cytometric analysis after staining with isotype control or anti-HLA-II mAb or anti-WT1 mAb.

**Supplementary Figure 8:**
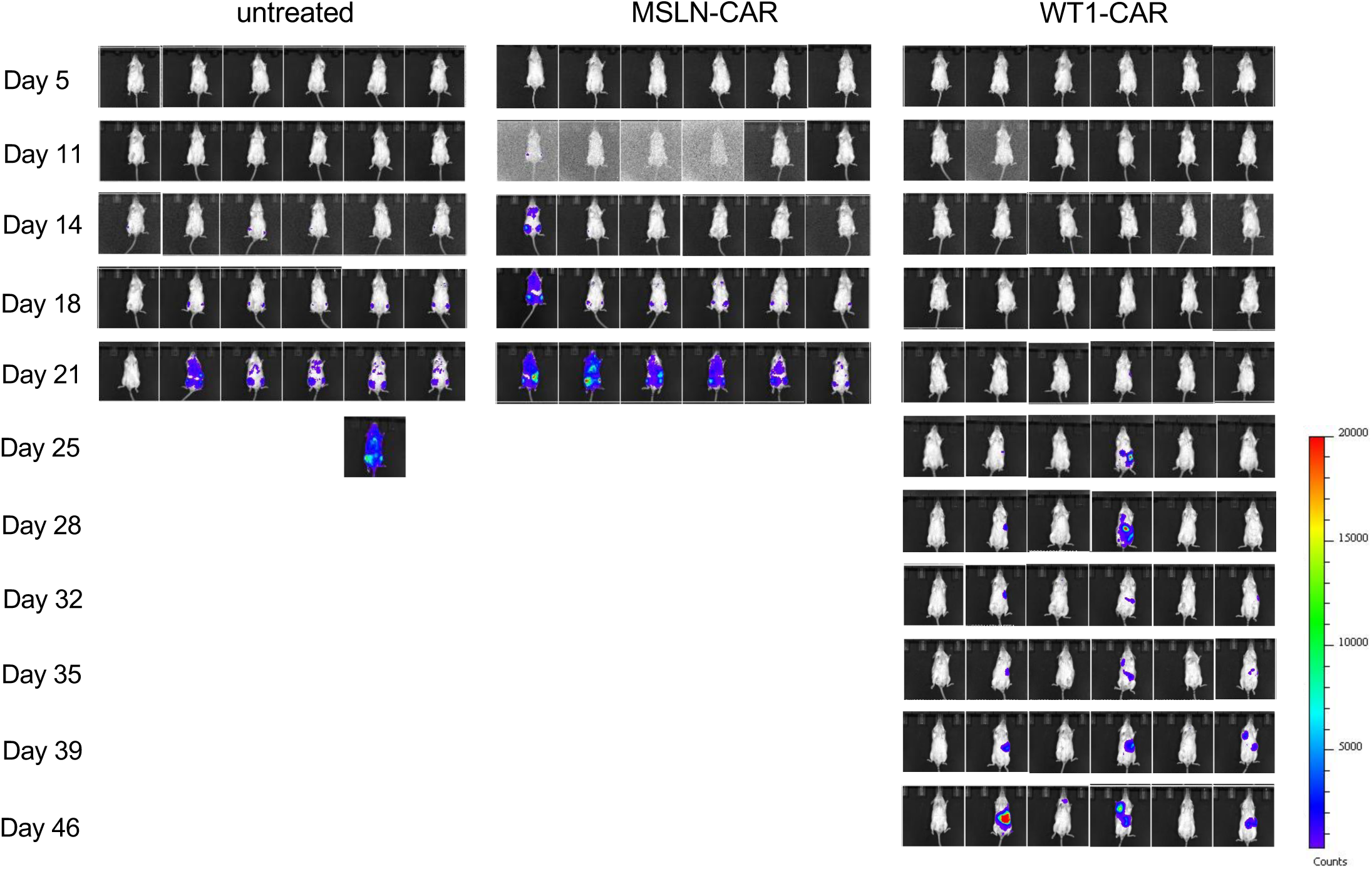
Antitumor activity of WT1-CAR T cells *in vivo*. *in vivo* bioluminescent imaging of luciferase activity in NSG mice in each treatment cohort.

**Supplementary Table S1.**
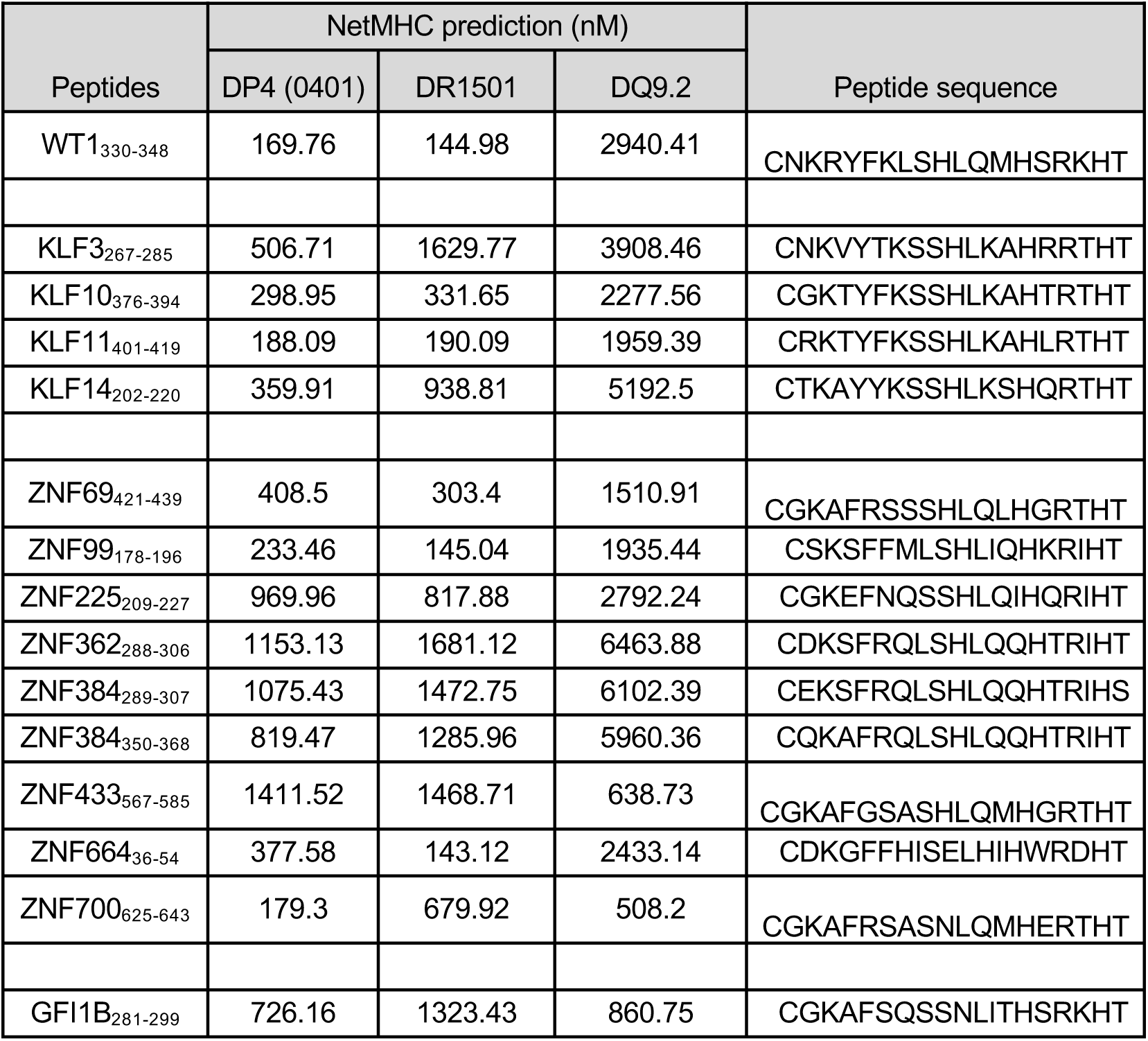

**Supplementary Table S2.**

| Cell line | Locus | Allele 1 | Allele 2 |
| --- | --- | --- | --- |
| HL60 | DPA1 | DPA1*01:03:01 | DPA1*02:01:01 |
|  | DPB1 | DPB1*04:01:01 | DPB1*13:01:01 |
|  | DQA1 | DQA1*02:01:01 | DQA1*02:01:01 |
|  | DQB1 | DQB1*03:03:02 | DQB1*03:03:02 |
|  | DRB1 | DRB1*07:01:01 | DRB1*07:01:01 |
|  | DRB345 | DRB4*01:03:01:02N | DRB4*01:03:01:02N |
| OCI-AML5 | DPA1 | DPA1*01:03:01 | DPA1*02:01:01 |
|  | DPB1 | DPB1*01:01:02 | DPB1*04:02:01 |
|  | DQA1 | DQA1*02:01:01 | DQA1*02:01:01 |
|  | DQB1 | DQB1*02:02:01 | DQB1*03:03:02 |
|  | DRB1 | DRB1*07:01:01 | DRB1*07:01:01 |
|  | DRB345 | DRB4*01:03:01 | DRB4*01:03:01:02N |
| U937/PR9 | DPA1 | DPA1*01:03:01 | DPA1*02:02:02 |
|  | DPB1 | DPB1*03:01:01 | DPB1*05:01:01 |
|  | DQA1 | DQA1*01:02:02 | DQA1*01:04:01 |
|  | DQB1 | DQB1*05:02:01 | DQB1*05:03:01 |
|  | DRB1 | DRB1*14:54:01 | DRB1*16:01:01 |
|  | DRB345 | DRB3*02:02:01 | DRB5*02:02:01 |

## Reference

1. Rosenberg SA, Restifo NP. Adoptive cell transfer as personalized immunotherapy for human cancer. Science. 2015;348:62–8.

2. June CH, Sadelain M. Chimeric Antigen Receptor Therapy. N Engl J Med. 2018;379:64–73.

3. Hsiue EH-C, Wright KM, Douglass J, Hwang MS, Mog BJ, Pearlman AH, et al. Targeting a neoantigen derived from a common TP53 mutation. Science. 2021;371:eabc8697.

4. Zajac P, Schultz-Thater E, Tornillo L, Sadowski C, Trella E, Mengus C, et al. MAGE-A Antigens and Cancer Immunotherapy. Front Med (Lausanne). 2017;4:18.

5. Raman MCC, Rizkallah PJ, Simmons R, Donnellan Z, Dukes J, Bossi G, et al. Direct molecular mimicry enables off-target cardiovascular toxicity by an enhanced affinity TCR designed for cancer immunotherapy. Sci Rep. 2016;6:18851.

6. Leung CSK. Endogenous antigen presentation of MHC class II epitopes through non-autophagic pathways. Front Immunol. 2015;6:464.

7. Roche PA, Furuta K. The ins and outs of MHC class II-mediated antigen processing and presentation. Nat Rev Immunol. 2015;15:203–16.

8. Yamashita Y, Anczurowski M, Nakatsugawa M, Tanaka M, Kagoya Y, Sinha A, et al. HLA-DP84Gly constitutively presents endogenous peptides generated by the class I antigen processing pathway. Nat Commun. 2017;8:15244.

9. Anczurowski M, Yamashita Y, Nakatsugawa M, Ochi T, Kagoya Y, Guo T, et al. Mechanisms underlying the lack of endogenous processing and CLIP-mediated binding of the invariant chain by HLA-DP84Gly. Sci Rep. 2018;8:4804.

10. Axelrod ML, Cook RS, Johnson DB, Balko JM. Biological Consequences of MHC-II Expression by Tumor Cells in Cancer. Clin Cancer Res. 2019;25:2392–402.

11. Kudela P, Janjic B, Fourcade J, Castelli F, Andrade P, Kirkwood JM, et al. Cross-Reactive CD4+ T Cells against One Immunodominant Tumor-Derived Epitope in Melanoma Patients. J Immunol. 2007;179:7932–40.

12. Galperin M, Farenc C, Mukhopadhyay M, Jayasinghe D, Decroos A, Benati D, et al. CD4+ T cell–mediated HLA class II cross-restriction in HIV controllers. Sci Immunol. 2018;3:eaat0687.

13. Laghmouchi A, Kester MGD, Hoogstraten C, Hageman L, de Klerk W, Huisman W, et al. Promiscuity of Peptides Presented in HLA-DP Molecules from Different Immunogenicity Groups Is Associated With T-Cell Cross-Reactivity. Front Immunol. 2022;13.

14. Denzin LK, Robbins NF, Carboy-Newcomb C, Cresswell P. Assembly and intracellular transport of HLA-DM and correction of the class II antigen-processing defect in T2 cells. Immunity. 1994;1:595–606.

15. Hou T, Macmillan H, Chen Z, Keech CL, Jin X, Sidney J, et al. An insertion mutant in DQA1*0501 restores susceptibility to HLA-DM: implications for disease associations. J Immunol. 2011;187:2442–52.

16. Fallang L-E, Bergseng E, Hotta K, Berg-Larsen A, Kim C-Y, Sollid LM. Differences in the risk of celiac disease associated with HLA-DQ2.5 or HLA-DQ2.2 are related to sustained gluten antigen presentation. Nat Immunol. 2009;10:1096–101.

17. Tawara I, Kageyama S, Miyahara Y, Fujiwara H, Nishida T, Akatsuka Y, et al. Safety and persistence of WT1-specific T-cell receptor gene−transduced lymphocytes in patients with AML and MDS. Blood. 2017;130:1985–94.

18. Chapuis AG, Egan DN, Bar M, Schmitt TM, McAfee MS, Paulson KG, et al. T cell receptor gene therapy targeting WT1 prevents acute myeloid leukemia relapse post-transplant. Nat Med. 2019;25:1064–72.

19. Fujiki F, Oka Y, Kawakatsu M, Tsuboi A, Nakajima H, Elisseeva OA, et al. A WT1 protein-derived, naturally processed 16-mer peptide, WT1332, is a promiscuous helper peptide for induction of WT1-specific Th1-type CD4+ T cells. Microbiol Immunol. 2008;52:591–600.

20. Anguille S, Fujiki F, Smits EL, Oji Y, Lion E, Oka Y, et al. Identification of a Wilms’ tumor 1-derived immunogenic CD4+ T-cell epitope that is recognized in the context of common Caucasian HLA-DR haplotypes. Leukemia. 2013;27:748–50.

21. Lin Y, Fujiki F, Katsuhara A, Oka Y, Tsuboi A, Aoyama N, et al. HLA-DPB1*05: 01-restricted WT1332-specific TCR-transduced CD4+ T lymphocytes display a helper activity for WT1-specific CTL induction and a cytotoxicity against leukemia cells. J Immunother. 2013;36:159–70.

22. Katsuhara A, Fujiki F, Aoyama N, Tanii S, Morimoto S, Oka Y, et al. Transduction of a Novel HLA-DRB1*04:05-restricted, WT1-specific TCR Gene into Human CD4+ T Cells Confers Killing Activity Against Human Leukemia Cells. Anticancer Res. 2015;35:1251–61.

23. Shafer P, Kelly LM, Hoyos V. Cancer Therapy With TCR-Engineered T Cells: Current Strategies, Challenges, and Prospects. Front Immunol. 2022;13:835762.

24. Campillo-Davo D, Flumens D, Lion E. The quest for the best: How TCR affinity, avidity, and functional avidity affect TCR-engineered T-cell antitumor responses. Cells. 2020;9:1720.

25. Stone JD, Chervin AS, Kranz DM. T-cell receptor binding affinities and kinetics: impact on T-cell activity and specificity. Immunology. 2009;126:165–76.

26. Fesnak AD, June CH, Levine BL. Engineered T cells: the promise and challenges of cancer immunotherapy. Nat Rev Cancer. 2016;16:566–81.

27. Abramson J, Adler J, Dunger J, Evans R, Green T, Pritzel A, et al. Accurate structure prediction of biomolecular interactions with AlphaFold 3. Nature. 2024;630:493–500.

28. Peyron I, Hartholt RB, Pedró-Cos L, van Alphen F, Brinke AT, Lardy N, et al. Comparative profiling of HLA-DR and HLA-DQ associated factor VIII peptides presented by monocyte-derived dendritic cells. Haematologica. 2018;103:172–8.

29. Karnaukhov V, Paes W, Woodhouse IB, Partridge T, Nicastri A, Brackenridge S, et al. HLA variants have different preferences to present proteins with specific molecular functions which are complemented in frequent haplotypes. Front Immunol. 2022;13.

30. Ariyaratana S, Loeb DM. The role of the Wilms tumour gene (WT1) in normal and malignant haematopoiesis. Expert Rev Mol Med. 2007;9:1–17.

31. Yarmarkovich M, Marshall QF, Warrington JM, Premaratne R, Farrel A, Groff D, et al. Targeting of intracellular oncoproteins with peptide-centric CARs. Nature. 2023;623:820– 7.

32. Ataie N, Xiang J, Cheng N, Brea EJ, Lu W, Scheinberg DA, et al. Structure of a TCR-Mimic Antibody with Target Predicts Pharmacogenetics. J Mol Biol. 2016;428:194–205.

33. Godkin AJ, Smith KJ, Willis A, Tejada-Simon MV, Zhang J, Elliott T, et al. Naturally Processed HLA Class II Peptides Reveal Highly Conserved Immunogenic Flanking Region Sequence Preferences That Reflect Antigen Processing Rather Than Peptide-MHC Interactions1. J Immunol. 2001;166:6720–7.

34. Painter CA, Stern LJ. Structural insights into HLA-DM mediated MHC II peptide exchange. Curr Top Biochem Res. 2011;13:39–55.

35. Sidney J, Steen A, Moore C, Ngo S, Chung J, Peters B, et al. Five HLA-DP molecules frequently expressed in the worldwide human population share a common HLA supertypic binding specificity. J Immunol. 2010;184:2492–503.

36. Davenport MP, Quinn CL, Chicz RM, Green BN, Willis AC, Lane WS, et al. Naturally processed peptides from two disease-resistance-associated HLA-DR13 alleles show related sequence motifs and the effects of the dimorphism at position 86 of the HLA-DR beta chain. Proc Natl Acad Sci U S A. 1995;92:6567–71.

37. Sant’Angelo DB, Robinson E, Janeway CA Jr, Denzin LK. Recognition of core and flanking amino acids of MHC class II-bound peptides by the T cell receptor. Eur J Immunol. 2002;32:2510–20.

38. Zavala-Ruiz Z, Strug I, Walker BD, Norris PJ, Stern LJ. A hairpin turn in a class II MHC-bound peptide orients residues outside the binding groove for T cell recognition. Proc Natl Acad Sci U S A. 2004;101:13279–84.

39. de Haan EC, Wauben MHM, Grosfeld-Stulemeyer MC, Kruijtzer JAW, Liskamp RMJ, Moret EE. Major histocompatibility complex class II binding characteristics of peptoid-peptide hybrids. Bioorg Med Chem. 2002;10:1939–45.

40. Ferrante A, Gorski J. Cooperativity of hydrophobic anchor interactions: evidence for epitope selection by MHC class II as a folding process. J Immunol. 2007;178:7181–9.

41. Garcia KC, Degano M, Teyton L, Wilson IA. Structural basis of TCR-pMHC recognition. J Neuroimmunol. 1998;90:5.

42. Tian J, Lehmann PV, Kaufman DL. T cell cross-reactivity between coxsackievirus and glutamate decarboxylase is associated with a murine diabetes susceptibility allele. J Exp Med. 1994;180:1979–84.

43. Kobayashi H, Song Y, Hoon DSB, Appella E, Celis E. Tumor-reactive T Helper Lymphocytes Recognize a Promiscuous MAGE-A3 Epitope Presented by Various Major Histocompatibility Complex Class II Alleles1. Cancer Res. 2001;61:4773–8.

44. Thomas R, Al-Khadairi G, Roelands J, Hendrickx W, Dermime S, Bedognetti D, et al. NY-ESO-1 Based Immunotherapy of Cancer: Current Perspectives. Front Immunol. 2018;9.

45. Lin X-X, Xie Y-M, Zhao S-J, Liu C-Y, Mor G, Liao A-H. Human leukocyte antigens: the unique expression in trophoblasts and their crosstalk with local immune cells. Int J Biol Sci. 2022;18:4043–52.

46. Yang S, Cohen CJ, Peng PD, Zhao Y, Cassard L, Yu Z, et al. Development of optimal bicistronic lentiviral vectors facilitates high-level TCR gene expression and robust tumor cell recognition. Gene Ther. 2008;15:1411–23.

47. Motmaen A, Dauparas J, Baek M, Abedi MH, Baker D, Bradley P. Peptide-binding specificity prediction using fine-tuned protein structure prediction networks. Proc Natl Acad Sci U S A. 2023;120:e2216697120.

48. Whitlow M, Bell BA, Feng SL, Filpula D, Hardman KD, Hubert SL, et al. An improved linker for single-chain Fv with reduced aggregation and enhanced proteolytic stability. Protein Eng. 1993;6:989–95.

49. Kochenderfer JN, Feldman SA, Zhao Y, Xu H, Black MA, Morgan RA, et al. Construction and Pre-clinical Evaluation of an Anti-CD19 Chimeric Antigen Receptor. J Immunother. 2009;32:689–702.

50. Carpenito C, Milone MC, Hassan R, Simonet JC, Lakhal M, Suhoski MM, et al. Control of large, established tumor xenografts with genetically retargeted human T cells containing CD28 and CD137 domains. Proc Natl Acad Sci U S A. 2009;106:3360–5.

51. Osborn MJ, Panoskaltsis-Mortari A, McElmurry RT, Bell SK, Vignali DAA, Ryan MD, et al. A picornaviral 2A-like sequence-based tricistronic vector allowing for high-level therapeutic gene expression coupled to a dual-reporter system. Mol Ther. 2005;12:569–74.

52. Hirano N, Butler MO, Xia Z, Ansén S, von Bergwelt-Baildon MS, Neuberg D, et al. Engagement of CD83 ligand induces prolonged expansion of CD8+ T cells and preferential enrichment for antigen specificity. Blood. 2006;107:1528–36.

53. Butler MO, Lee J-S, Ansén S, Neuberg D, Hodi FS, Murray AP, et al. Long-Lived Antitumor CD8+ Lymphocytes for Adoptive Therapy Generated Using an Artificial Antigen-Presenting Cell. Clin Cancer Res. 2007;13:1857–67.

54. Sugata K, Matsunaga Y, Yamashita Y, Nakatsugawa M, Guo T, Halabelian L, et al. Affinity-matured HLA class II dimers for robust staining of antigen-specific CD4+ T cells. Nat Biotechnol. 2021;39:958–67.

55. Butler MO, Imataki O, Yamashita Y, Tanaka M, Ansén S, Berezovskaya A, et al. Ex Vivo Expansion of Human CD8+ T Cells Using Autologous CD4+ T Cell Help. PLoS One. 2012;7:e30229.

56. Zheng Z, Chinnasamy N, Morgan RA. Protein L: a novel reagent for the detection of Chimeric Antigen Receptor (CAR) expression by flow cytometry. J Transl Med. 2012;10:29.

57. Gadalla R, Boukhaled GM, Brooks DG, Wang BX. Mass cytometry immunostaining protocol for multiplexing clinical samples. STAR Protoc. 2022;3:101643.

